# Monocyte Subsets with High Osteoclastogenic Potential and Their Epigenetic Regulation Orchestrated by IRF8

**DOI:** 10.1101/2020.06.02.126284

**Authors:** Amitabh Das, Xiaobei Wang, Jessica Kang, Alyssa Coulter, Amol C. Shetty, Mahesh Bachu, Stephen R. Brooks, Stefania Dell’Orso, Brian L. Foster, Xiaoxuan Fan, Keiko Ozato, Martha J. Somerman, Vivek Thumbigere-Math

**Author notes:** Correspondence (V.T.M).

## Abstract

Osteoclasts (OCs) are bone resorbing cells formed by the serial fusion of monocytes. In mice and humans, three distinct subsets of monocytes exist; however, it is unclear if all of them exhibit osteoclastogenic potential. Here we show that in wild-type mice, Ly6C^hi^ and Ly6C^int^ monocytes are the primary source of OC formation when compared to Ly6C^−^ monocytes. Their osteoclastogenic potential is dictated by increased expression of signaling receptors and activation of pre-established transcripts, as well as de novo gain in enhancer activity and promoter changes. In the absence of IRF8, a transcription factor important for myelopoiesis and osteoclastogenesis, all three monocyte subsets are programmed to display higher osteoclastogenic potential. Enhanced NFATc1 nuclear translocation and amplified transcriptomic and epigenetic changes initiated at early developmental stages direct the increased osteoclastogenesis in *Irf8* deficient mice. Collectively, our study provides novel insights into the transcription factors and active *cis*-regulatory elements that regulate OC differentiation.

## INTRODUCTION

Osteoclasts (OCs) are multinucleated giant cells that resorb bone and play an important role in the development and remodeling of skeleton. OCs that control fetal skeletal development and tooth eruption originate in fetal ossification centers from embryonic erythroid-myeloid progenitors (EMP) (Jacome-Galarza et al., 2019). In adults, OC maintenance and function are facilitated by serial fusion of hematopoietic stem cell (HSC)-derived circulating monocytes with long-lived OC syncytia (Jacome-Galarza et al., 2019).

Monocytes develop in the bone marrow (BM) from MDP-cMOP axis (monocyte and dendritic cell progenitors-common monocyte progenitors), and are subsequently released into circulation (Geissmann et al., 2010; Ginhoux and Jung, 2014; Guilliams et al., 2018; Hettinger et al., 2013). Under the influence of microenvironmental cues, monocytes get recruited to peripheral tissues where they differentiate into a variety of cells including OCs, macrophages, and dendritic cells (DCs) (Xing et al., 2005). In humans and mice, functionally two distinct subsets of monocytes exist (Geissmann et al., 2003; Passlick et al., 1989). The “inflammatory or classical monocytes” represent Ly6C^hi^ monocytes in mice and CD14^hi^ CD16^−^ monocytes in humans, which can differentiate into M1 macrophages or TipDCs to produce pro-inflammatory cytokines and exhibit antimicrobial activities (Dominguez and Ardavin, 2010; Guilliams et al., 2018; Nahrendorf et al., 2007; Satoh et al., 2017; Serbina and Pamer, 2006). In contrast, “nonclassical or patrolling monocytes” represent Ly6C^−^ monocytes in mice and CD14^low^ CD16^hi^ monocytes in humans (Auffray et al., 2007; Carlin et al., 2013), which survey the luminal surfaces of blood vessels for dead cells and can differentiate into M2 macrophages to orchestrate tissue repair. Recent transcriptomic studies have highlighted that heterogeneity exists in mice and human monocytes and have identified a third intermediate population (Ly6C^int^ and CD14^hi^ CD16^low^, respectively) (Mildner et al., 2017; Villani et al., 2017; Wong et al., 2011). However, the precise function of this intermediate population is unknown.

Development of Ly6C^hi^ monocytes is tightly regulated by lineage determining transcription factors (TFs) such as *Irf8* (Kurotaki et al., 2013), *Spi1* (PU.1) (DeKoter et al., 1998; McKercher et al., 1996), *Ccr2* (Serbina and Pamer, 2006), and *Klf4* (Alder et al., 2008; Kurotaki et al., 2013). Whereas, the generation of Ly6C^−^ monocytes is controlled by *Nr4a1* (Nur77) (Hanna et al., 2011; Thomas et al., 2016) and C/EBPβ (Mildner et al., 2017; Tamura et al., 2017). The development of Ly6C^int^ monocytes remains unknown.

Currently, it is unclear if all monocyte subsets exhibit osteoclastogenic potential. To date, limited studies have explored this question and the results are conflicting (Ammari et al., 2018; Charles et al., 2012; Jacome-Galarza et al., 2013; Misharin et al., 2014; Park-Min et al., 2013; Puchner et al., 2018; Seeling et al., 2013; Zhao et al., 2015). Furthermore, the role of lineage determining TFs in regulating OC differentiation on a genome-wide scale remains poorly understood. IRF8, apart from playing an important role in myelopoiesis (Hambleton et al., 2011; Kurotaki et al., 2019; Kurotaki et al., 2013; Tamura et al., 2000; Tamura and Ozato, 2002), also functions as a negative regulator of OC by inhibiting NFATc1, a master regulator of osteoclastogenesis (Fang et al., 2016; Thumbigere-Math et al., 2019; Zhao et al., 2009). In this study, we utilized a novel *Irf8* conditional knockout (*Irf8 cKO*) mouse model to characterize the importance of IRF8 in OC progenitor development and epigenetic regulation of OC differentiation. We show that *Irf8 cKO* mice contain severely diminished Ly6C^hi^ monocytes but retain their ability to generate Ly6C^−^ monocytes, and all three monocyte subsets in *Irf8 cKO* mice exhibit increased potential for OC formation. In contrast, Ly6C^hi^ and Ly6C^int^ monocytes are the main source of OC formation in wild-type (WT) mice. Ly6C^hi^ and Ly6C^int^ monocytes were found to express higher levels of RANK and CCR2, indicating that these cells constitute a primed source of pre-OCs. RNA-seq data overlapped with ChIP-seq data showed that the differentiation of Ly6C^hi^ and Ly6C^int^ monocytes into OCs ensured significant transcriptomic changes specific to each subset, which was accompanied by de novo gain in enhancer activity and promoter changes. In *Irf8 cKO* mice, the transcriptomic and epigenetic changes are amplified and changes occurring during the progenitor stage may be critical for initiating early lineage commitment and priming monocyte subsets to differentiate into robust OCs. Collectively, our study provides critical insights into the complex network of TFs and active *cis*-regulatory elements that drive OC differentiation process in WT and *Irf8 cKO* mice.

## RESULTS

### Generation of *Irf8 cKO* Mice and Characterization of HSCs

Previously, IRF8’s role in osteoclastogenesis has been studied using *Irf8* global knock-out (*Irf8 gKO*) mice (Izawa et al., 2019; Thumbigere-Math et al., 2019; Zhao et al., 2009). However, the severely altered population and properties of HSCs in *Irf8 gKO* mice (Holtschke et al., 1996; Tamura and Ozato, 2002), along with the development of chronic myelogenous leukemia and splenomegaly, has limited the genome-wide analysis of IRF8 function in OC precursors derived from a monocyte/macrophage lineage. Hence to overcome this limitation, we generated *Irf8 cKO* mice (*Irf8*^*fl/fl*^; *Csf1r*^*cre*^) by crossing *Irf8*^*fl/fl*^ mice (Feng et al., 2011) with *Csf1r*^*cre*^ mice (Deng et al., 2010) (Figure S1A-H). Deletion of IRF8 was confirmed by the absence of IRF8 transcript and protein expression in BM monocytes (BMMs) of *Irf8 cKO* mice (Figure S1E-F). The development of HSCs in *Irf8 cKO* mice was examined by flow cytometric analysis by comparing BM, blood, and spleen cells for expression of cell lineage markers. We observed that *Irf8 cKO* mice accumulate MDPs and cMoPs and have decreased monocyte and increased neutrophil population (Figures 1A-B and S2A-B). IRF8 is known to enforce monocyte development by opposing neutrophil lineage (Becker et al., 2012; Kurotaki et al., 2014). In the absence of IRF8, myeloid progenitors accumulate at the MDP and cMoP stages and fail to generate their downstream population, instead aberrantly giving rise to neutrophils (Kurotaki et al., 2014; Sichien et al., 2016). In contrast to *Irf8 gKO* mice, we found that cMoPs and monocytes in *Irf8 cKO* mice were less severely affected (Figures 1A and S2A).

**Figure 1.**
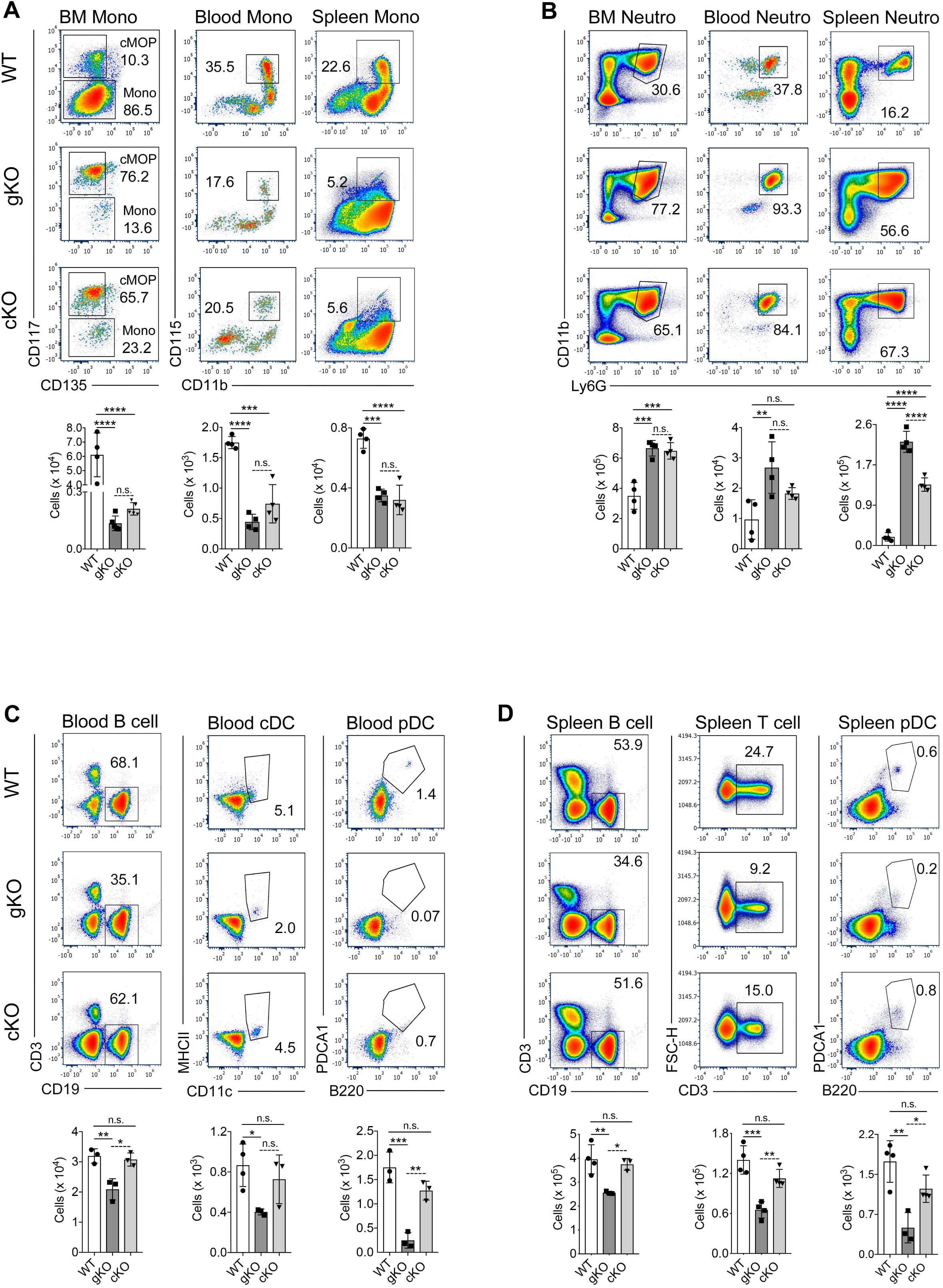
Characterization of HSCs in *Irf8 cKO* Mice. (Cossarizza et al.) Flow cytometric analysis of major immune cells in BM, blood, and spleen. Pseudocolor plots show cell population in percentages and bar graphs show absolute counts. Data are representative of at least four independent experiments (n = 4-5 mice per genotype). See also Figure S2 for extensive flow cytometric analysis of HSCs. Cells were gated as described in Supplemental Methods.

Additionally, IRF8 promotes the differentiation of common lymphoid progenitors (CLPs) to B cells (Shin et al., 2011; Wang et al., 2019; Wang et al., 2008), drives effector differentiation of CD8 T cells (Miyagawa et al., 2012), and regulates the development of type 1 conventional DCs (cDC1s) as well as plasmacytoid DCs (pDCs) (Aliberti et al., 2003; Becker et al., 2012; Kurotaki et al., 2019; Schiavoni et al., 2002; Tsujimura et al., 2003). Consequently, *Irf8 gKO* mice have reduced B- and T-cells, completely lack cDC1s and pDCs, and are immunodeficient. In contrast to *Irf8 gKO* mice, we found that the development of B- and T-lymphocytes, cDCs, and pDCs were less severely affected in *Irf8 cKO* mice BM, blood, and spleen (Figures 1C-D and S2C-F).

### *Irf8 cKO* Mice Exhibit Increased Osteoclastogenesis and Provide Novel Osteoclast Transcriptomic Results

IRF8 deficiency in OC precursors but not osteoblasts is known to promote osteoclastogenesis (Thumbigere-Math et al., 2019; Zhao et al., 2009). Hence, to study the effects of conditional deletion of *Irf8* in OC precursors, we analyzed *Irf8 cKO* mice for abnormal bone and OC phenotypes. We noted that *Irf8 cKO* mice display severe osteoporosis accompanied by dramatic decreases in total amount of bone (39%), trabecular number (35%), and trabecular BMD (35%) when compared to WT mice (Figure 2A). Histological analysis showed a substantial increase in OC numbers and resorption activity in *Irf8 cKO* mice (Figure 2B). Increased OC-mediated bone resorption was supported by ∼2.5-fold increase in serum C-terminal telopeptides type I collagen (CTX) levels in *Irf8 cKO* mice (Figure 2C).

**Figure 2.**
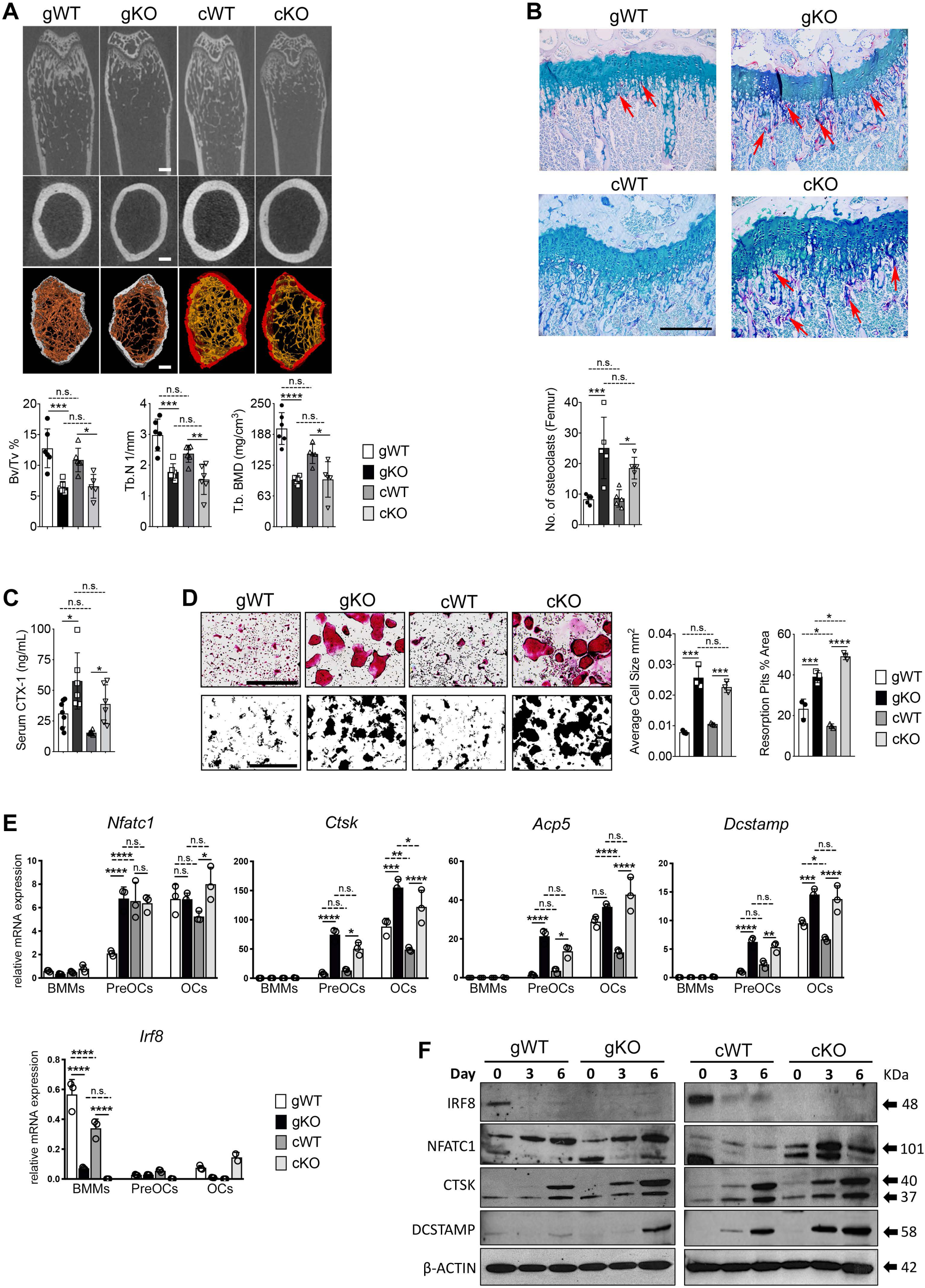
*Irf8 cKO* Mice Exhibit Increased Osteoclastogenesis. (A) Micro-CT analysis of femurs (8.5-week-old mice). Top, longitudinal view; middle, axial view of the cortical bone in midshaft; bottom, axial view of the trabecular bone in metaphysis. Scale bar, 0.5 mm. Bar graphs show bone morphometric analysis of femurs. BV/TV, bone volume per tissue volume; Tb.N, trabecular number; Tb. BMD, trabecular bone mineral density. n = 5-6 mice per genotype. (B) Histological analysis of proximal femurs from 8.5-week-old mice (TRAP-stained; red arrows indicate OCs). Scale bar, 1000 μm. n = 5 mice per genotype. (C) Serum CTX-1 levels were measured in 8.5 to 9.5-week-old mice. n = 6-7 mice per genotype. (D) Top panel, TRAP-stained cells show OC formation. Bottom panel shows resorption activity of OCs. Scale bar, 1000 μm. Bar graphs show quantified number of average cell size of TRAP^+^ cells and percentage area of resorption in each group. (E) mRNA expression of OC-specific genes in BMMs, Pre-OCs, and OCs, as measured by RT-qPCR. (F) Immunoblot analysis of OC-specific proteins. Day0=BMMs, Day3=PreOCs, and Day6=OCs. (D-F) Data are representative of three independent experiments performed in triplicates. (G) Error bars indicate mean ± STD. \**p* < 0.05, \*\**p* < 0.01, \*\*\**p* < 0.001, \*\*\*\**p* < 0.0001, and n.s. = non-significant. See also Figure S3 for novel osteoclast transcriptomic results in *Irf8 cKO* Mice. See Table S1 for qPCR primer sequences.

To measure OC formation in vitro, we cultured BMMs from WT and *Irf8 cKO* mice with M□CSF and RANKL for 6 days. Significantly increased number of OCs formed in *Irf8 cKO* cultures with cell size ∼2-fold larger than WT cells, which was appended by a 3-fold increase in resorption activity (Figure 2D). Correspondingly, the mRNA and protein expression of NFATc1 and its downstream OC-related genes were strongly upregulated in *Irf8 cKO* OCs when compared to WT OCs (Figure 2E-F). Taken together, these results establish the loss of IRF8 regulatory function in *Irf8 cKO* mice, and *Irf8 cKO* mice present OC and skeletal phenotypes similar to *Irf8 gKO* mice.

Next, we aimed to understand if less altered HSCs or conditional deletion of *Irf8* in monocyte/macrophage lineage provide a better model for studying genomewide effects of IRF8 on OC transcriptome. We performed RNA-seq on *Irf8* WT, *cKO*, and *gKO* BMMs, preosteoclasts (PreOCs), and OCs (Figure S3A). 4726 RANKL-responsive genes were found to be commonly regulated among all groups (Figure S3B) and they clustered into four major categories with different enriched functions (Figure S3C). Transcripts enriched in *Irf8 cKO* mice were associated with hematopoietic cell differentiation, interferon signaling, chemokine receptor binding, and inflammatory response, suggesting that these functions are operational in *Irf8 cKO* mice when compared to *Irf8 gKO* mice (Figure S3D). We noted that a greater number of genes were significantly upregulated in *Irf8 cKO* OCs when compared to WT and *Irf8 gKO* OCs (Figure S3E-G). Several novel RANKL-responsive genes were identified in *Irf8 cKO* OCs that were diminished in *Irf8 gKO* OCs (61% vs 17%, Figure S3F), which included genes such as *Ctsg, Eps8, Prr11, Mpo, Myb, Kif23*, and *Ccnb2*. GSEA analyses comparing cell-specific transcript signatures obtained from Ingenuity Pathway Analysis (IPA) further illustrated that *Irf8 cKO* mice provide more precise/reliable OC-specific outcomes when compared to *Irf8 gKO* mice (Figure S3H). Altogether, these findings indicate that *Irf8 cKO* is an improved model over *Irf8 gKO* to study OCs, and it provides novel OC transcriptomic results that are deficient in *Irf8 gKO* mice.

### All Monocyte Subsets in *Irf8 cKO* Mice Exhibit Increased Potential for Osteoclast Formation

While the total number of monocytes were diminished in *Irf8 cKO* mice (Figures 1A and S2A), the animals exhibited increased osteoclastogenesis (Figures 2 and S3). This raises the question whether all monocyte subsets are equally diminished in *Irf8 cKO* mice or if particular monocyte subset development is relatively maintained, and if those subsets have increased potency for OC formation.

Flow cytometric analyses identified that Ly6C^hi^ monocytes were significantly diminished in blood, spleen, and BM of *Irf8 cKO* mice when compared with WT mice (Figure 3A). Ly6C^int^ monocytes were also significantly reduced in blood and BM of *Irf8 cKO* mice, but not in spleen. The counts of Ly6C^−^ monocytes were reduced in BM of *Irf8 cKO* mice, however their numbers in blood and spleen were not significantly different than WT mice.

**Figure 3.**
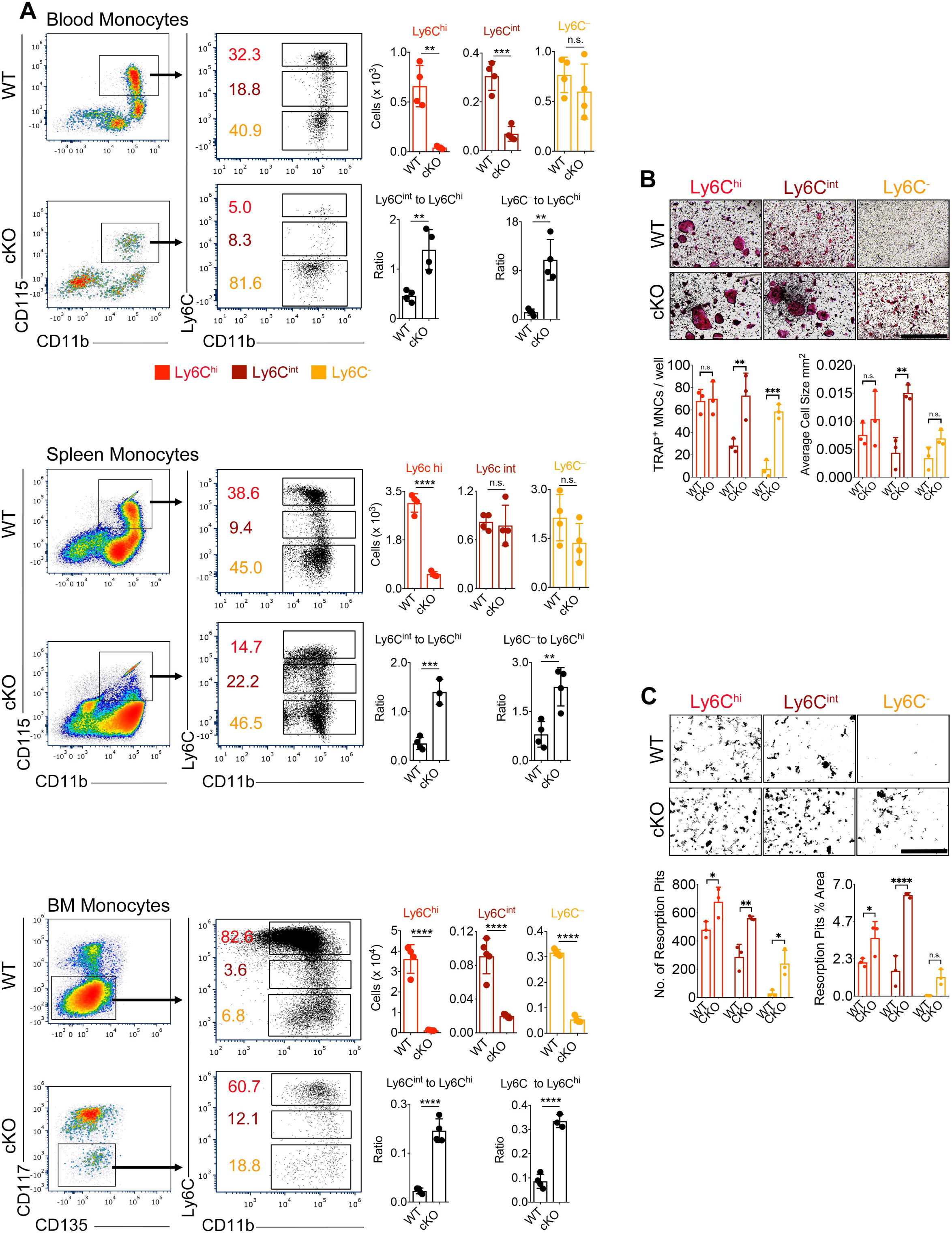
*Irf8 cKO* Mice Retain Development of Circulating Ly6C^−^ Monocytes and All Monocyte Subsets in *Irf8 cKO* Mice Exhibit Increased Potential for Osteoclast Formation. (A) Flow cytometric analysis of monocyte subsets in blood, spleen, and BM of WT and *Irf8 cKO* mice. Pseudocolor plots show cell population in percentages and bar graphs show absolute counts. Data are representative of at least four independent experiments (n = 4-5 mice per genotype). (B) TRAP-stained cells show OC formation potential of monocyte subsets. Data are mean of three independent experiments performed in triplicates. Scale bar, 1000 μm. (C) Resorption activity of OCs. Data are representative of three independent experiments performed in triplicates. Scale bar, 1000 μm. (D) Error bars indicate mean ± STD. \*\**p* < 0.01, \*\*\**p* < 0.001, \*\*\*\**p* < 0.0001, and n.s. = non-significant.

Normally in WT mice, ∼85% of BM monocytes are Ly6C^hi^, 6% are Ly6C^int^, and 9% are Ly6C^−^ cells (Kurotaki et al., 2013; Mildner et al., 2016; Thomas et al., 2016). In blood, ∼35% of monocytes are Ly6C^hi^, 20% are Ly6C^int^, and 45% are Ly6C^−^ cells. In comparison, we observed that in *Irf8 cKO* mice the ratios of Ly6C^−^ and Ly6C^int^ monocytes to Ly6C^hi^ monocytes were significantly higher (Figure 3A). In *Irf8 cKO* mice, increased number of OCs can be detected as early as 15 days of age (data not shown). We found that the ratios of Ly6C^−^ and Ly6C^int^ monocytes to Ly6C^hi^ monocytes were significantly higher at 15 days of age in *Irf8 cKO* BM and blood and remained elevated at every time point measured (15 days, 5 wks, 8.5 wks, 12 wks, data not shown). Altogether, these results indicate that *Irf8 cKO* mice retain their ability to generate Ly6C^−^ monocytes, but at a reduced efficiency, and most of the circulating monocytes in *Irf8 cKO* mice are Ly6C^−^ cells.

Next, we investigated the osteoclastogenic potential of Ly6C^hi^, Ly6C^int^, and Ly6C^−^ monocytes in WT and *Irf8 cKO* mice. Ly6C^hi^, Ly6C^int^, and Ly6C^−^ BM monocytes were isolated by FACS and cultured with M-CSF and RANKL. The sorted monocyte subsets exhibited varying phenotypic plasticity and differentiated into mature OCs much sooner than unsorted cells (4 days vs 6 days). In WT mice, Ly6C^hi^ and Ly6C^int^ but not Ly6C^−^ monocytes exhibited increased potential for OC formation and resorption activity (Figure 3B-C). In stark contrast, all monocyte subsets in *Irf8 cKO* mice robustly differentiated into mature OCs and the differences against respective WT subsets were-Ly6C^hi^ = 1-fold, Ly6C^int^ = 2-fold, and Ly6C^−^ = 6-fold (Figure 3B). Accordingly, Ly6C^hi^ (1-fold), Ly6C^int^ (2-fold), and Ly6C^−^ (9-fold) OCs from *Irf8 cKO* mice demonstrated significantly increased resorption activity when compared to respective WT subsets (Figure 3C). Altogether, these findings imply that IRF8 deficiency not only affects the development of selective monocyte subsets, but also promotes the osteoclastogenic potential of all three monocyte subsets.

### Molecular Mechanisms Regulating the Osteoclastogenesis Potential of Monocyte Subsets in WT and *Irf8 cKO* mice

To understand the plausible mechanisms shaping the varying osteoclastogenic potential of Ly6C^hi^, Ly6C^int^, and Ly6C^−^ monocytes in WT and *Irf8 cKO* mice, we examined the expression of cell surface receptors involved in OC signaling. We assessed the expression of RANK, CCR2, and CX_3_CR1 in BMMs by flow cytometry. In both WT and *Irf8 cKO* mice, we noted that Ly6C^hi^ monocytes expressed high levels of RANK, Ly6C^int^ monocytes expressed moderate levels of RANK, and Ly6C^−^ monocytes expressed low levels of RANK (Figure 4A). The expression of CCR2 followed a pattern similar to RANK in both WT and *Irf8 cKO* subsets. The expression of CX_3_CR1 was inverse to RANK and CCR2. These results indicate that the expression level of RANK, CCR2, and CX_3_CR1 may regulate the activation of OC signaling pathways and OC formation potential of distinct monocyte subsets. Indeed, in WT mice, Ly6C^hi^ monocytes that express high levels of RANK and CCR2 differentiated into robust OCs, Ly6C^int^ monocytes that express moderate levels of RANK and CCR2 formed moderate OCs, and Ly6C^−^ monocytes that express low levels of RANK and CCR2 formed minimal OCs (Figure 3B-C). These findings were corroborated by mRNA and protein expression of OC-related genes that correlated with the osteoclastogenic potential of Ly6C^hi^, Ly6C^int^, and Ly6C^−^ subsets (Figure 4B-C).

**Figure 4.**
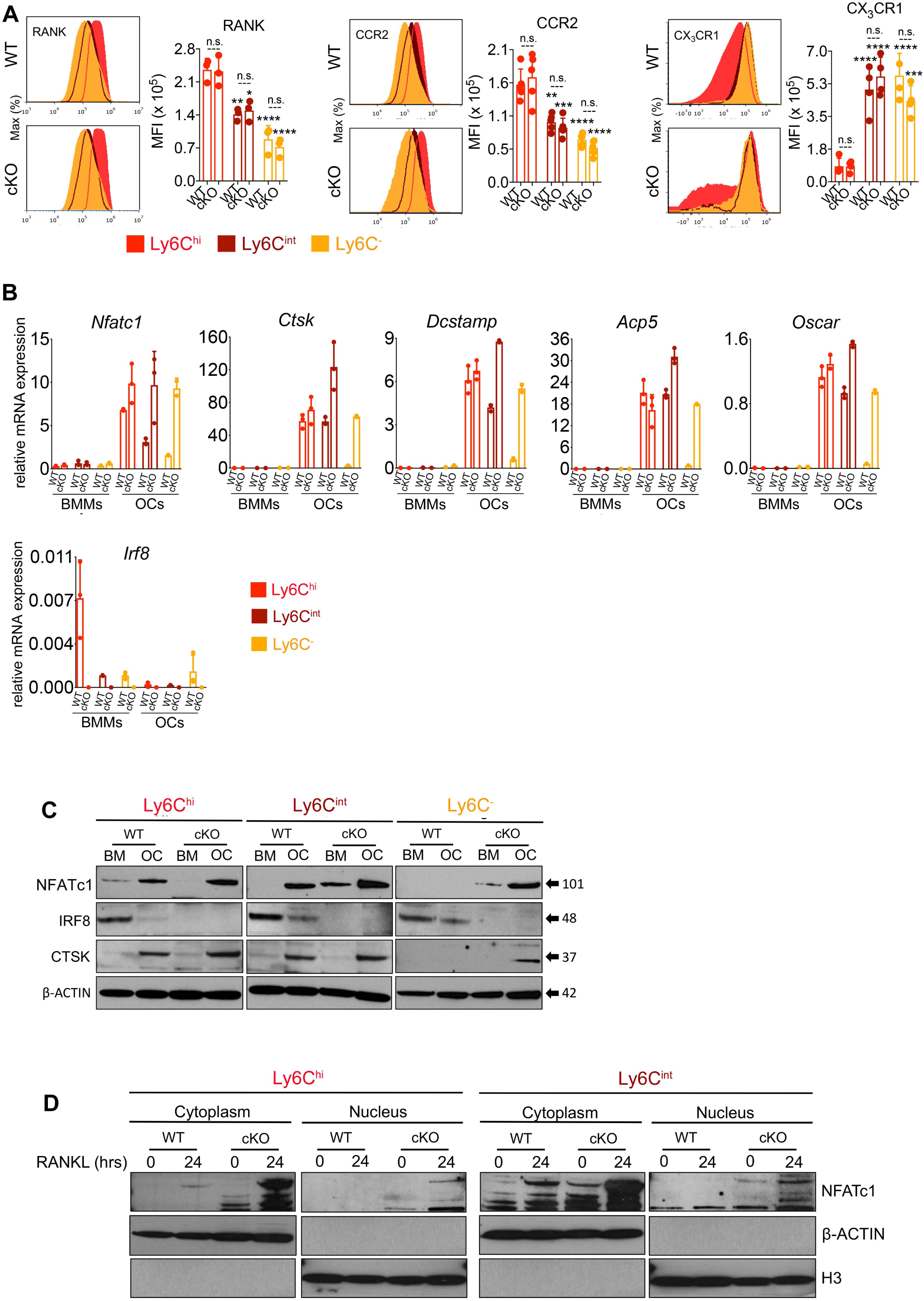
Osteoclastogenic Potential of Monocyte Subsets is Regulated by Cell Surface Receptors and Increased Nuclear Translocation of NFATc1 in *Irf8 cKO* Mice. (A) Histograms display the expression of cell surface markers in monocyte subsets (without RANKL). Bar graphs show median ± IQR frequencies of at least three independent experiments. \**p* < 0.05, \*\**p* < 0.01, \*\*\**p* < 0.001, \*\*\*\**p* < 0.0001, and n.s. = non-significant. (B) mRNA expression of OC-specific genes in distinct subsets. (C) Immunoblot analysis of OC-specific proteins in distinct subsets. (D) Immunoblot analysis of cytoplasmic and nuclear NFATc1 expression at 0 hours and 24 hours after RANKL stimulation. See also Table S1 for qPCR primer sequences.

While the expression of RANK, CCR2, and CX_3_CR1 in *Irf8 cKO* subsets were similar to WT mice, comparatively all monocyte subsets in *Irf8 cKO* mice formed increased OCs and the mRNA and protein expression of OC-related genes were strongly upregulated in *Irf8 cKO* OCs when compared to WT OCs. In depth analysis indicated that in *Irf8 cKO* mice, there is increased nuclear translocation of NFATc1 within 24 hours of RANKL stimulation (Figure 4D). The IAD domain of IRF8 is known to physically interact with the TAD-A domain of NFATc1 and inhibit NFATc1 nuclear translocation (Jiang et al., 2014). In *Irf8 cKO* mice, the early release of NFATc1 from inhibition by IRF8 may enable higher auto-amplification of NFATc1, thus resulting in strong induction of NFATc1 expression and enhanced activation of downstream target genes that promote OC formation. Collectively, these results suggest that the osteoclastogenic potential of monocyte subsets may be regulated by the expression level of OC signaling receptors, and further influenced by IRF8-mediated NFATc1 nuclear translocation.

### Transcriptional Program Governing the Osteoclastogenic Potential of Ly6C^hi^, Ly6C^int^, and Ly6C^−^ Monocytes Subsets in WT and *Irf8 cKO* mice

To identify global transcriptional program governing the osteoclastogenic potential of Ly6C^hi^, Ly6C^int^, and Ly6C^−^ monocytes, and how it may be further influenced by IRF8 deficiency, we performed RNA-seq on sorted Ly6C^hi^, Ly6C^int^, and Ly6C^−^ cells from WT and *Irf8 cKO* mice. Cells were analyzed prior to (BMMs) and 4 days after RANKL stimulation (OCs) (Figure S4A). During the course of OC differentiation, 4833 RANKL-responsive genes (FDR <0.01, >2-fold difference in any pairwise comparison among a total of 38,942 genes) were commonly regulated between WT and *Irf8 cKO* subsets (Figure S4B). Hierarchical clustering classified the genes into four major categories with different enriched functions (Figure S4B-C). Cluster I consisted of genes primarily upregulated in WT BMMs and were associated with type 1 IFN signaling, cytokine signaling, and innate immune response. Cluster II comprised of genes co-expressed by WT and *Irf8 cKO* BMMs, and were correlated with cellular response to cytokine stimulus, extracellular matrix organization, and neutrophil mediated immunity. Cluster III was defined by genes predominantly upregulated in *Irf8 cKO* BMMs and were associated with DNA replication and cell cycle pathway. Cluster IV was characterized by genes upregulated in both WT and *Irf8 cKO* OCs and comprised of genes involved in OC regulation such as *Nfatc1, Ctsk, Oscar, Dcstamp, Acp5, Dnmt3a*, and *Calcr*, etc. Cluster IV gene expression was prominent in Ly6C^hi^ and Ly6C^int^ OCs in both genotypes, moderate in *Irf8 cKO* Ly6C^−^ OCs, and diminished in WT Ly6C^−^ OCs.

Amenable to its name as inflammatory monocytes, Ly6C^hi^ BMMs exhibited distinct inflammatory signature when compared to Ly6C^−^ BMMs, but also displayed augmented genes important for OC differentiation (Figure S4D). Similar findings were noted in Ly6C^int^ BMMs. These results indicate that the developmentally established transcriptome of Ly6C^hi^ and Ly6C^int^ monocytes is likely a feature of committed OC precursors, thus predisposing them to respond intensely to RANKL stimulation (Figures S5A-B). This notion was further apparent when RANKL-stimulated Ly6C^hi^, Ly6C^int^, and Ly6C^−^ OCs were compared against their respective BMMs (Figure S5C). In WT mice, Ly6C^hi^ OCs were markedly enriched for genes involved in OC differentiation and oxidative phosphorylation, Ly6C^int^ OCs were enriched for genes involved in DNA replication and RNA transport, and Ly6C^−^ OCs were enriched for genes involved in chemokine signaling and cell adhesion (Figure S5D). In contrast, all three subset OCs in *Irf8 cKO* mice were overrepresented by genes involved in OC signaling, such as MAPK and PI3K-Akt signaling in Ly6C^hi^, TCA cycle and pyruvate metabolism in Ly6C^int^, and PI3K-Akt-mTOR and EGFR1 signaling in Ly6C^−^ (Figure S5D).

Subsequent direct comparison of *Irf8 cKO* vs WT subsets identified a greater number of genes important for OC differentiation were significantly regulated in all three subsets of *Irf8 cKO* BMMs and OCs (Figure 5A-B). Notably, in *Irf8 cKO* Ly6C^hi^, Ly6C^int^, and Ly6C^−^ BMMs, the basal expression of many established positive regulators of osteoclastogenesis were strongly enriched and negative regulators were diminished (Figure 5A). These results imply that genetic ablation of *Irf8* initiates the OC differentiation program and lineage commitment of all three monocyte subsets at a very early stage. Consistent with BMMs, *Irf8 cKO* subset OCs further strongly expressed several positive regulators (e.g., *Nfatc1, Acp5, Ctsk, Calcr, Mmp9, Dcstamp*) of OC differentiation when compared to WT OCs (Figure 5A, C). Many of the differentially expressed genes (DEGs) contained IRF8 binding sites, suggesting that they may be direct transcriptional targets of IRF8.

**Figure 5.**
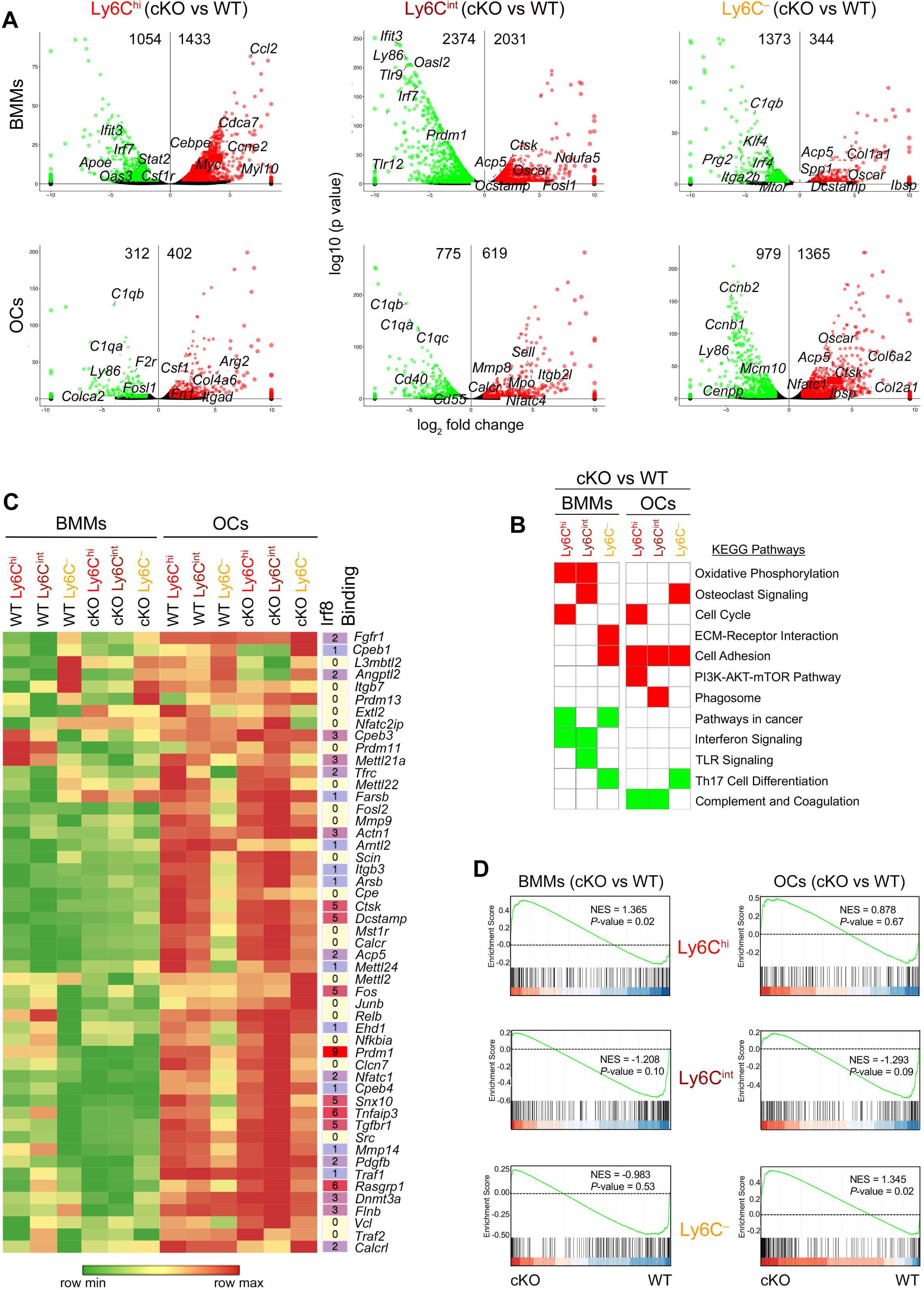
Transcriptional Profiling of Ly6C^hi^, Ly6C^int^, and Ly6C^−^ BMMs and OCs in *Irf8 cKO* Mice. (A) Volcano plot of transcriptomic changes between *Irf8 cKO* vs WT subsets (one-way ANOVA, >2-fold change, FDR<0.01). red=up, green=down, back=no significant change. (B) Heatmap showing GO term enrichment for genes in each cluster. (C) Gene expression changes for randomly selected osteoclast-specific markers (n=51). Shaded heat map on the right indicates the presence of one or more IRF8 binding sites. (D) GSEA analysis of osteoclast-specific signatures in *Irf8 cKO* vs WT subsets. See also Figures S4-S5 for detailed transcriptomic analysis.

Further, to investigate the effect of *Irf8* deficiency on ontogeny and activity of OCs and OC precursors, we performed GSEA analysis, which showed strong enrichment of OC-specific signatures in *Irf8 cKO* Ly6C^hi^, Ly6C^int^, and Ly6C^−^ BMMs and OCs when compared to WT cells (Figure 5D). Collectively, these results corroborate the findings noted in Figure 4 and demonstrate that WT Ly6C^hi^ and Ly6C^int^ monocytes developmentally contain OC-specific transcripts, which are augmented upon RANKL stimulation. Additionally, IRF8 deficiency initiates the OC differentiation program and lineage commitment at a very early stage. In *Irf8 cKO* mice, OC-specific transcripts in all three monocyte subsets are primed to robustly respond to RANKL stimulation and various OC signaling pathways are activated in each subset.

### Induction of Active *cis-*Regulatory Elements Critical for Osteoclast Differentiation

Currently, the epigenetic mechanisms governing the OC differentiation process remains poorly understood. To investigate the combinatorial activity of TFs, *cis*-regulatory elements, and the enhancer landscape that establish and maintain OC transcriptional identity, we profiled histone modifications (H3K4me1, H3K4me3, H3K27ac, H3K27me3) and PU.1 and IRF8 binding by ChIP-seq. RNA-seq data from sorted Ly6C^hi^, Ly6C^int^, and Ly6C^−^ cells were overlaid with ChIP-seq data to identify epigenetic changes specific to each subset and genotype. Active promoters were identified according to proximity to transcription start sites (TSS) (<2 kb) and H3K4me3, while active enhancers were defined by their distance from TSS (>2 kb) and enrichment of both H3K4me1 and H3K27ac (Creyghton et al., 2010; Heintzman et al., 2007). Transcriptional repressors were identified by their distance from TSS (<2 kb) and enrichment of H3K27me3.

RNA-seq data indicated that majority (88%-90%) of the RANKL-responsive genes in WT monocytes had IRF8 occupancy (Figure S6A). Analysis of H3K4me3 marks for these IRF8 dependent genes (Figure S6B) identified several promoters that were either established (13%-24%) or lost (30%-37%) during the course of OC differentiation in WT Ly6C^hi^, Ly6C^int^, and Ly6C^−^ OCs (Figure S6C). Among them, 2-5% were active promoters that were established de novo (Figure S6D) and included OC-specific genes such as Ly6C^hi^ - *Ccr1, Myo1e, Angpt2, Prr11, Mt1*; Ly6C^int^ - *Ccr1, Angpt2, Prr11, Cenpf, Mt1*; and Ly6C^−^ - *Mmp9, Mmp14, Myo1b, Mefv, Creg2*. Subsequently, when H3K4me3 signal was analyzed for DEGs between *Irf8 cKO* vs WT mice (Figure 6A), we observed that *Irf8 cKO* BMMs (10%-38%) and OCs (17%-21%) had gained increased promoters compared to their respective WT cells (Figure 6B). Of the established promoters, 4-5% were active promoters specific to *Irf8 cKO* BMMs (Figure 6C) and included OC-related genes such as Ly6C^hi^ - *Elane, Trem3, Serpinb1a, Ly6c2, Myb, Rasgrp3*; Ly6C^int^ - *Elane, Phgdh, Rasgrp3, Dmd, Trem3, Myb*; and Ly6C^−^ - *Rasgrp3, Siglec-E, Clec12a*. Likewise, 6-10% were active promoters specific to *Irf8 cKO* OCs (Figure 6C) and included genes such as Ly6C^hi^ - *Dkk2, Cd4, Ffar4, Ltb, Cd69*; Ly6C^int^ - *Dkk2, Mpo, Elane, Camp, Mmp8*, Siglec-E; and Ly6C^−^ - *Myl9, Itga11, Dlx3, Il34, Mmp8*.

**Figure 6.**
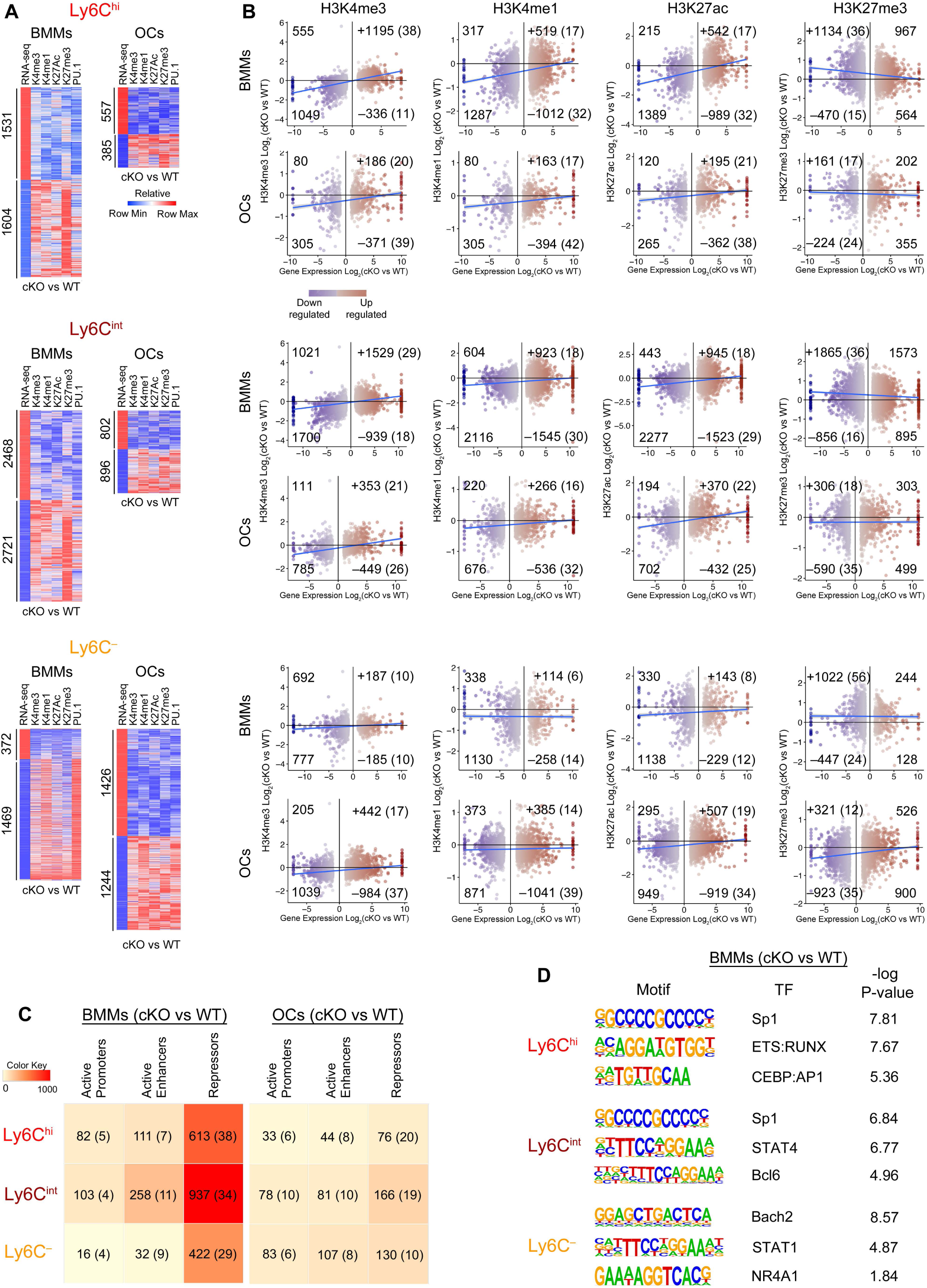
Histone Modifications Regulated by IRF8 Deficiency During Osteoclastogenesis. (A) Heatmap illustrating H3K4me3, H3K4me1, H3K27ac, H3K27me3, and PU.1 binding signal in DEGs between *Irf8 cKO* vs WT BMMs and OCs. (B) Scatter plots show the correlation between histone modifications and genes either upregulated or downregulated in *Irf8 cKO* vs WT BMMs and OCs. Gain in histone marks indicated by + and loss of histone marks indicated by –. Data are represented as log_2_. (C) Heatmap depicting number of active promoters, active enhancers, and repressors acquired in *Irf8 cKO* vs WT BMMs and OCs. Numbers in parenthesis indicate percentages. (D) Homer motif analysis of active promoter and active enhancer regions enriched in *Irf8 cKO* vs WT BMMs. See also Figure S6 for additional histone modification data.

Analysis of H3K4me1 marks for IRF8-dependent DEGs identified de novo enhancer acquisitions in WT subset OCs (19%-31%) (Figure S6C). These gained enhancers were located in the vicinity of genes such as *Nfatc1, Calcr, Ctsk, Oscar, Dcstamp, Dnmt3a, Vegfc*, and *Relb*, which positively regulate OCs (Figures 7A and S7A). During BMM to OC differentiation, 23%-31% of the genes lost enhancer marks (Figure S6C) and they included mainly negative regulators of OCs such as *Bcl6, Bcl2, Mafb, Irf4, Irf7, Cx3cr1, Nr4a1*, and *Klf6* (Figure S7B). Subsequently, when H3K4me1 signal was analyzed for DEGs between *Irf8 cKO* vs WT mice, we noted that *Irf8 cKO* BMMs and OCs had either acquired (BMMs=6%-18%, OCs=14%-17%) or lost (BMMs=14%-32%, OCs=32%-42%) more enhancers when compared to WT cells (Figure 6B). The gained enhancers in BMMs were located in the vicinity of OC-related genes such as *Spp1, Atp6v0d2, Prr11, Mpo, Angpt2, Ccl3, Mmp12*, and *Msr1*. The gained enhancers in OCs were located in the vicinity of genes such as *Csf1, Calcr, Trem3, Ccr2, Il1r2, Atp6v0d2, Nfkbia, Cxcr2, Notch1*, and *Rasgrp3*. To validate these results, we analyzed H3K27ac signal in WT and *Irf8 cKO* subsets. H3K27ac signal in enhancers followed a pattern similar to H3K4me1 (Figures S6C and 6B), corroborating that the establishment of enhancers in the vicinity of OC-specific genes is essential for enrichment of OC transcription (Figures 7A and S7A). Prominent changes were evident when H3K4me1 and H3K27ac marks were overlapped to identify active enhancers. Active enhancers were increasingly noted at BMM stage in *Irf8 cKO* Ly6C^hi^ and Ly6C^int^ cells, and in contrast noted at OC stage in *Irf8 cKO* Ly6C^−^ monocytes (Figure 6C). These active enhancer findings match with transcriptomic results (Figure 5A), indicating that in the absence of IRF8, enhancer establishment occurring during the progenitor stage may be critical for priming monocytes to differentiate into robust OCs, which is supported by the increased OC phenotype noted in Ly6C^hi^ and Ly6C^int^ cells when compared to Ly6C^−^ monocytes.

**Figure 7.**
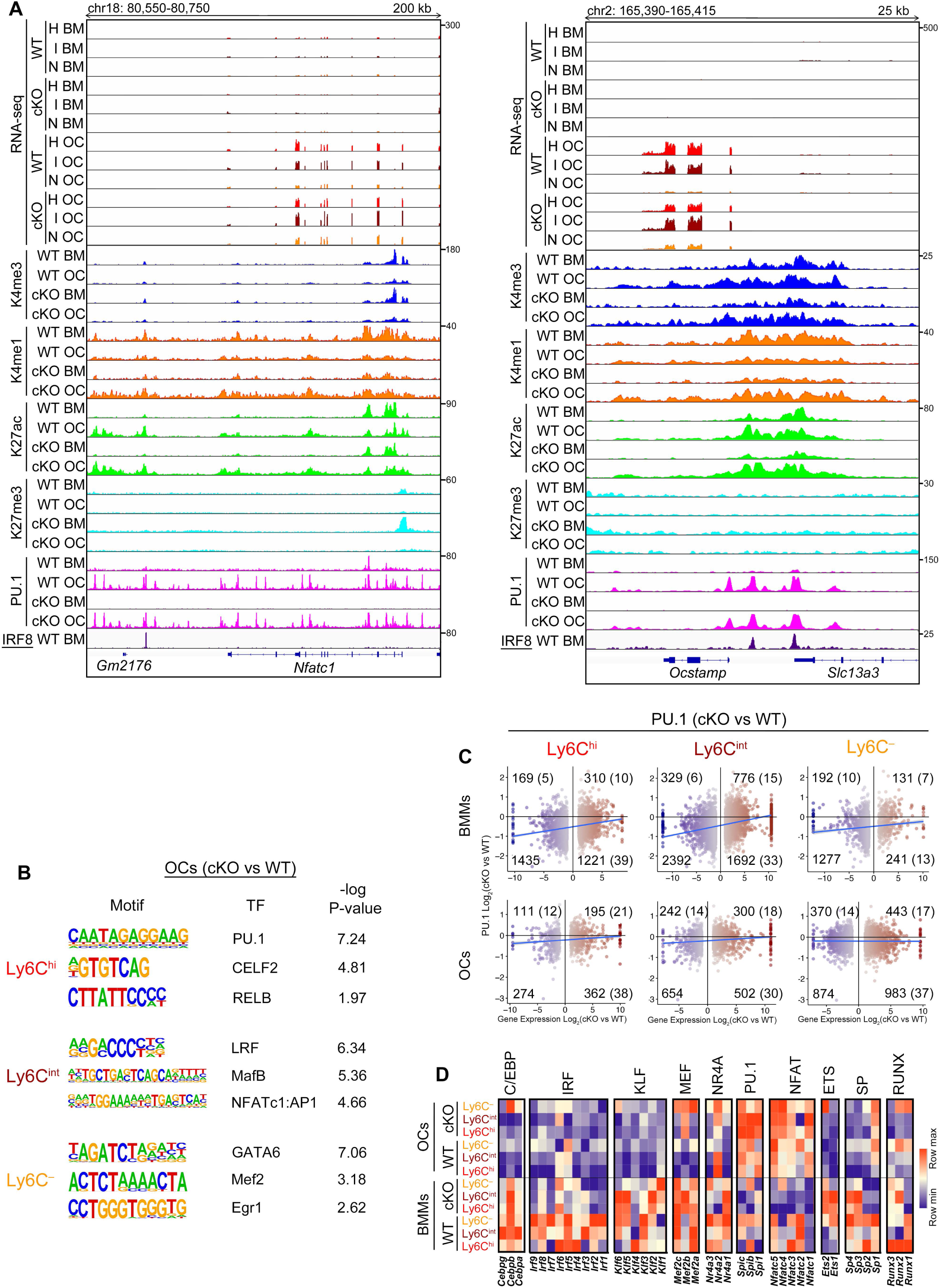
The Epigenetic Landscape of WT and *Irf8 cKO* BMMs and OCs. (A) UCSC Genome Browser tracks showing normalized tag-density profiles at key OC-specific genes. Note the enrichment of H3K4me1 and H3K27ac marks at the *Nfatc1* and *Ocstamp* loci in *Irf8 cKO* OCs when compared to WT OCs. (B) Homer motif analysis of active promoter and active enhancer regions enriched in *Irf8 cKO* vs WT OCs. (C) Scatter plots show the correlation between PU.1 binding and genes either upregulated or downregulated in *Irf8 cKO* vs WT BMMs and OCs. Data are represented as log_2_. (D) RNA-seq expression of TFs predicted to bind motifs identified in Figures 6D, 7B and S6E. See also Figures S7 for examples of IGV tracks.

While H3K27ac is associated with active transcription, deposition of H3K27me3 by the polycomb repressive complex (PRC)2 is associated with chromatin-based gene silencing (Di Croce and Helin, 2013). To examine if IRF8 regulates OC-specific genes by methylation of H3K27 around the TSS, we assessed H3K27me3 marks. A global increase in H3K27me3 intensity was observed among the downregulated genes in both WT and *Irf8 cKO* subsets (Figure S6C-D and Figure 6B-C). Among the positive regulators of OCs, H3K27me3 intensity was significantly decreased at the OC stage when compared to their respective BMMs stage (Figures 7A and S7A). The concordant loss of H3K27me3 was associated with gain in promoters and enhancers and enrichment of transcription. In contrast, among the negative regulators of OCs, H3K27me3 intensity was significantly increased at the OC stage, which was associated with transcriptional repression (Figure S7B). All of these differences were stark in *Irf8 cKO* BMMs and OCs when compared to WT cells.

To identify potential TFs responsible for transcriptome and epigenome regulation during OC differentiation, we performed motif analysis within the identified active promoter and active enhancer regions. We identified both previously known and novel candidate regulators in WT and *Irf8 cKO* Ly6C^hi^, Ly6C^int^, or Ly6C^−^ BMMs and OCs (Figures S6E, 6D and 7B). Most of these TFs have known functions in OC differentiation, and are potential targets of IRF8 and PU.1. Previously, it has been reported that during osteoclastogenesis, PU.1 switches its transcription partner from IRF8 to NFATc1 and alters the chromatin binding regions, which is associated with changes in epigenetic profiles and gene expression (Izawa et al., 2019). Therefore, we investigated the relationship between IRF8 and PU.1 binding sites in BMMs and OCs. Among the IRF8-dependent DEGs in WT subsets, 90% of them contained PU.1 binding sites (Figure S6C), highlighting the significant overlap between IRF8 and PU.1 binding sites in BMMs. The majority of PU.1 binding regions were shared between BMMs and OCs. Among the DEGs between *Irf8 cKO* vs WT mice, only a small fraction displayed enhanced PU.1 binding sites in *Irf8 cKO* subsets (Figure 7C). PU.1 peaks in OCs were enriched for histone marks in both WT and *Irf8 cKO* subsets. Figures 7A and S7 are representative of ChIP-seq data showing the co-localization of PU.1 with IRF8 and promoter, enhancer, and repressor marks in BMMs and OCs.

Subsequently, using RNA-seq data we compared predicted motif enrichment with TF expression in Ly6C^hi^, Ly6C^int^, and Ly6C^−^ subsets (Figure 7D). PU.1 and NFAT family of TFs were prominently expressed in OCs, confirming the requirement of these TFs for OC differentiation. In contrast, IRF8 and KLF family of TFs, which are known negative regulators of OC differentiation were decreased. Altogether, these data suggest that the Ly6C^hi^ and Ly6C^int^ monocytes are transcriptionally and epigenetically programmed to differentiate in mature OCs when compared to Ly6C^−^ anti-inflammatory monocytes. A combinatorial activity of TFs and *cis*-regulatory elements in the vicinity of several OC-specific genes are responsible for establishing and maintaining OC identity in these subsets. In *Irf8 cKO* mice, several promoters and enhancers established during the progenitor stage may be critical for initiating early lineage commitment and priming monocyte subsets to differentiate into robust OCs. In *Irf8 cKO* mice, while amplified transcriptomic and epigenetic changes are noted in well-known OC-specific genes, the stark differences against WT OCs are more visible in the lesser known OC-specific genes.

## DISCUSSION

OCs are the only known bone resorbing cells in the body. Defective OC function leads to osteopetrosis (Tolar et al., 2004), whereas, excessive OC activity leads to bone loss and osteoporosis (Boyle et al., 2003). Understanding the molecular mechanisms that control OC development and function is vital for developing effective intervention and therapeutic strategies for OC-related disorders.

Previous attempts to gain an in-depth understanding of IRF8’s role in osteoclastogenesis have been limited by the extremely altered HSCs in *Irf8 gKO* mice (Holtschke et al., 1996; Tamura and Ozato, 2002). The *Irf8 cKO* mice generated in this study provide an improved mouse model as they exhibit less severely altered HSCs, but present OC and skeletal phenotypes similar to *Irf8 gKO* mice. Furthermore, the *Irf8 cKO* mice provide more precise and novel OC transcriptomic results when compared to *Irf8 gKO* mice. These results are in stark contrast to a previous study, which failed to notice in vivo skeletal or OC phenotypes in *Irf8*^*fl/fl*^; *Lyz2*^*cre*^ mice, but observed increased TRAP^+^ OCs in vitro (Saito et al., 2017). It is unclear if the *Irf8*^*fl/fl*^; *Lyz2*^*cre*^ mice in their study harbored extremely altered HSCs. These differences compared to our study could be related to different “macrophage targeting” Cre lines used and the efficacy and specificity in cell deletion (Abram et al., 2014). While our findings illustrate effective deletion of *Irf8* in the monocyte/macrophage lineage of *Irf8 cKO* mice, the possibility that minor non-specific targeting of other immune cells could have occurred cannot be ruled out.

During myelopoiesis, IRF8 is sharply expressed at the MDP stage, and IRF8 interacts with C/EBPα to restrain MDPs and cMoPs from differentiating into neutrophils (Kurotaki et al., 2014). IRF8 in turn binds to KLF4 to govern the differentiation of MDPs and cMoPs into monocytes (Kurotaki et al., 2013). Consistent with the loss of IRF8 function, we noted that *Irf8 cKO* mice accumulate myeloid progenitors at cMoP stage and fail to effectively generate downstream monocytes, instead aberrantly giving rise to neutrophils. In-depth analysis indicated that Ly6C^hi^ monocytes were severely diminished in *Irf8 cKO* mice, whereas, the generation of Ly6C^−^ monocytes was relatively maintained. Developmentally, Ly6C^−^ monocytes are considered to be derived from Ly6C^hi^ monocytes (Patel et al., 2017; Sugimoto et al., 2015; Varol et al., 2007; Yona et al., 2013). However, under certain circumstances Ly6C^−^ monocytes can arise directly from MDPs or cMoPs independently of Ly6C^hi^ monocytes (Hettinger et al., 2013; Yanez et al., 2017; Zhu et al., 2016). Perhaps, this explains the existence of Ly6C^−^ monocytes in *Irf8 cKO* mice despite the absence of Ly6C^hi^ monocytes. The existence of Ly6C^−^ monocytes could be further explained by the fact that IRF8 is more essential for the development of Ly6C^+^ cells than Ly6C^−^ monocytes (Kurotaki et al., 2013).

As monocytes contribute to OC formation, it is expected that all monocyte subsets would exhibit osteoclastogenic potential. However, to date, only few studies have explored this subject and the results are conflicting. The osteoclastogenic potential of Ly6C^int^ monocytes has never been studied. Some groups have shown that in BM, CD11b^−/low^ Ly6C^hi^ progenitors (i.e., cMoPs) rather than CD11b^+^ Ly6C^hi^ monocytes exhibit increased potential for OC formation (Charles et al., 2012; Jacome-Galarza et al., 2013). This is supported by the fact that CD11b and β2-integrin signaling negatively regulates OC differentiation by transiently repressing RANK expression and *Nfatc1* transcription (Park-Min et al., 2013). In contrast, other studies have shown that CD11b^+^ Ly6C^hi^ monocytes in BM and blood are far more efficient than Ly6C^−^ monocytes in differentiating into mature OCs (Ammari et al., 2018; Seeling et al., 2013; Zhao et al., 2015). In humans, it is firmly established that CD14^hi^ CD16^−^ classical monocytes but not nonclassical monocytes are the main source of OC formation (Komano et al., 2006; Sprangers et al., 2016). In agreement with the latter group of findings, we explicitly demonstrate that CD11b^+^ Ly6C^hi^ and Ly6C^int^ monocytes in WT mice are transcriptionally and epigenetically programmed to differentiate in mature OCs when compared to Ly6C^−^ monocytes. It is unclear if Ly6C^hi^, Ly6C^int^, and Ly6C^−^ monocytes exhibit varying osteoclastogenic potential in vitro vs in vivo, and in physiologic vs pathologic conditions. In inflammatory arthritis in mice, depending upon the experimental models used, previous studies have reported that either Ly6C^−^ (Misharin et al., 2014; Puchner et al., 2018) or Ly6C^hi^ (Ammari et al., 2018; Seeling et al., 2013) monocytes specifically migrate to the inflamed joints and contribute to bone erosion. Identifying specific subsets driving tissue destruction in a disease state is extremely critical for developing targeted anti-OC therapy.

Mechanistically, our findings suggest that the osteoclastogenic potential of monocyte subsets is regulated by OC signaling receptors such as RANK, CCR2 and CX_3_CR1. The binding of RANK receptor to its ligand RANKL triggers distinct signaling cascades and induces TFs such as NFATc1, c-Fos, and NF-κB to promote OC formation (Boyce and Xing, 2008; Boyle et al., 2003). Activation of CCR2 in OC precursors leads to increased RANK expression and predisposes cells to differentiate into mature OCs (Binder et al., 2009). CX_3_CR1 negatively correlate with Ly6C expression (Mildner et al., 2016) and favors the maintenance of osteoclastic precursors, but not differentiated osteoclasts (Hoshino et al., 2013; Koizumi et al., 2009). It has been reported that increased expression of RANK is likely a feature of committed OC precursors, and human monocytes expressing high levels of RANK are predisposed to produce two-fold increased OCs when compared to cells expressing moderate or low levels of RANK (Atkins et al., 2006). Consistent with these findings, we noted that Ly6C^hi^ monocytes that expressed higher levels of RANK and CCR2 formed increased OCs when compared to Ly6C^int^ and Ly6C^−^ monocytes in WT mice. Further in support of CCR2 influencing the osteoclastogenic potential of Ly6C^+^ monocytes, in vivo delivery of anti-CCR2 antibody (MC21) has been shown to selectively deplete Ly6C^hi^ monocytes and inhibit OC formation and bone destruction (Seeling et al., 2013). CX_3_CR1^−^ OCs are known to exhibit higher inflammatory bone resorption (Madel et al., 2020), which is in agreement with reduced CX_3_CR1 noted in Ly6C^hi^ subset. Collectively, our mechanistic studies provide critical insight into why Ly6C^hi^ and Ly6C^int^, but not Ly6C^−^ monocytes exhibit osteoclastogenesis potential, and how it may be further augmented by increased NFATc1 nuclear translocation in the absence of IRF8.

Normally, TFs bind to promoters near TSS and distal enhancers to regulate chromatin signature and gene expression (Heintzman et al., 2007; Heinz et al., 2015). Studying the states of active promoters and active enhancers is critical for understanding how gene expression patterns are established by TFs during cell differentiation. To date, the genomewide mapping of active *cis*-regulatory elements involved in OC differentiation has been very limited (Carey et al., 2018; Carey et al., 2019; Izawa et al., 2019; Rohatgi et al., 2018). In the present study, by overlapping RNA-seq data with ChIP-seq data, we identified *cis*-regulatory regions that are active during OC differentiation and are specific to each subset and genotype. During the course of BMMs to OC differentiation, several promoters and enhancers were either established or lost in WT and *Irf8 cKO* cells. The distribution and enrichment of promoters and enhancers were highly specific to the differentiation stage and varied between WT and *Irf8 cKO* BMMs and OCs. The de novo promoter and enhancer acquisitions in the vicinity of OC-specific genes integrated with H3K27me3-mediated transcriptional repression is extremely critical for establishing and maintaining OC transcriptional identity. We explicitly demonstrate that in a steady state, Ly6C^hi^ and Ly6C^int^ monocytes are the OC forming cells and their osteoclastogenic potential is dictated by activation of pre-established transcripts, as well as de novo gain in enhancer activity and promoter changes. IRF8 deficiency further augments these transcriptomic and epigenetic changes in *Irf8 cKO* Ly6C^hi^ and Ly6C^int^ subsets at an early developmental stage, priming them to display higher osteoclastogenic potential when compared to their WT counterparts. Ly6C^−^ cells do not display osteoclastogenic potential in WT mice, however, in the absence of IRF8, active enhancers and transcriptional enrichment are established later on during the course of OC differentiation that shape their OC formation potential in *Irf8 cKO* mice. In *Irf8 cKO* mice, the gain in enhancers and promoters were noted in well-known OC-specific genes, however, the differences against WT cells were more visible in the lesser known OC-specific genes. These lesser known OC-specific genes are the unique features of *Irf8 cKO* when compared to *Irf8 gKO* mice.

DNA motif analysis identified several known and novel transcriptional regulators in WT and *Irf8 cKO* BMMs and OCs. Most of these TFs have known functions in OC differentiation, and are potential targets of IRF8 and PU.1. While the majority of PU.1 binding regions were shared between BMMs and OCs, it has been reported that PU.1 heterodimerizes with IRF8 to regulate BMM development (Kurotaki et al., 2018), and switches its partner to MITF and NFATc1 to control OC differentiation (Carey et al., 2018; Izawa et al., 2019). Similarly, we noted that among the DEGs in Ly6C^hi^, Ly6C^int^, and Ly6C^−^ BMMs, there was a significant overlap between IRF8 and PU.1 binding sites. It is anticipated that these identified PU.1 binding regions will be overlapped with NFATc1 binding sites in OCs. Despite repeated attempts, we were unable to obtain successful NFATc1 ChIP-seq results due to the lack of commercially available ChIP-grade NFATc1 antibody. Nevertheless, our results are in agreement with previous observations by Izawa et al, which showed that RANKL induced downregulation of *Irf8* and upregulation of *Nfatc1* is associated with significant histone modifications (Izawa et al., 2019). Comparatively, the study by Izawa et al. had few shortcomings as ChIP-seq experiments did not include *Irf8* deficient mice (except for H3K27ac) or appropriate input controls. Our study overcomes these shortcomings and provides significant new information about the important epigenetic and transcriptomic changes occurring at early developmental stages that prime the *Irf8 cKO* progenitors to differentiate into robust OCs.

In summary, to the best of our knowledge, this is the first study to investigate the osteoclastogenic potential of monocyte subsets in a novel *Irf8 cKO* mice and characterize the epigenetic mechanisms governing OC differentiation process in distinct monocyte subsets. Our study is a shift from the conventional approach of studying monocytes as a whole population and provides evidence of OC heterogeneity. Our study demonstrates the need for OC research to focus on individual monocyte/OC subsets in order to develop targeted therapy for OC-related disorders. Finally, this study provides critical insights into the important role of IRF8 in osteoclastogenesis, and effective inhibition of IRF8 by molecules that target chromatin regulators may offer promising therapeutic approaches.

## Supporting information

Key Resource Table

## ACKNOWLEDGEMENTS

We thank Gustavo Gutierrez-Cruz and Faiza Naz of NIAMS genomic core facility for assistance with next-generation sequencing and data acquisition. We thank Kan Jiang of NIAMS genomic core facility for assistance with ChIP-seq data quality assessment. We thank Anup Mahurkar from Institute for Genome Sciences, University of Maryland School of Medicine for critical input into ChIP-seq analysis. We thank Michael C. Ostrowski and Sudarshana M. Sharma from Medical University of South Carolina for providing us PU.1 antibody. We thank Bryan Hahn and Karen Underwood from University of Maryland School of Medicine Flow Cytometry Shared Service for assistance with cell sorting. We thank Satish Yesupatham from University of Maryland School of Dentistry for assistance with FACS. We thank Kim C. Mansky from University of Minnesota for critically reviewing this manuscript. Funding sources: R00DE028439 to V.T.M.; start-up funds from Univ. of Maryland School of Dentistry to V.T.M.; University of Maryland Baltimore Institute of Clinical & Translational Research (ICTR) grant to V.T.M; R01DE027639 to B.L.F., and intramural funding to M.J.S. from NIAMS/NIH, and to K.O. from NICHD/NIH.

## AUTHOR CONTRIBUTIONS

V.T.M., A.D., and X.W., conceived the project and designed the experiments; A.D., X.W., V.T.M., and J.K., performed the experiments; A.D., performed and S.D. helped with RNA-seq and ChIP-seq data acquisition; X.F., advised on flow cytometry analysis and cell sorting; V.T.M., A.D., A.C.S., S.B., S.D., and M.B., performed computational analyses for RNA-seq and ChIP-seq; A.C., X.W., and B.L.F., performed micro-CT and histological analysis; M.J.S., and K.O., advised on experiments and provided intellectual input; V.T.M., wrote the paper, supervised the project, and secured funding; all authors contributed to reviewing and editing.

## DECLARATION OF INTERESTS

The authors declare no conflict of interests.

## STAR METHODS

### LEAD CONTACT

Further information and requests for resources and reagents should be directed to and will be fulfilled by the Lead Contact, Vivek Thumbigere-Math.

### MATERIALS AVAILABILITY

Mouse line generated in this study are available from the Lead Contact upon request with a completed Materials Transfer Agreement.

### DATA AND CODE AVAILABILITY

The accession numbers for the RNA-seq and ChIP-seq datasets reported in this paper can be found at GEO: GSE151483.

### EXPERIMENTAL MODEL AND SUBJECT DETAILS

The following mouse strains were used in this study: *Irf8*^*fl/fl*^ mice (Feng et al., 2011), *Csf1r*^*cre*^ mice (Deng et al., 2010), *Irf8* gKO mice and *Irf8* gWT mice (Tamura and Ozato, 2002; Thumbigere-Math et al., 2019). *Irf8 cKO* mice were generated by crossing *Irf8*^*fl/fl*^ mice with *Csf1r*^*cre*^ mice, a monocyte/macrophage targeting Cre driver mice. The resulting *Irf8 cWT* (*Irf8*^*fl*^; *Csf1r*^*cre/-*^) and *Irf8 cKO* (*Irf8*^*fl/fl*^; *Csf1r*^*cre/+*^) mice were both on C57BL/6 and mixed background, and mice from both backgrounds were utilized in this study with no significant differences noted in experimental results. Animals aged between 8-12 weeks were used for all experiments as noted in the figure legends. Additionally, animals aged between 15 days-12 weeks were used to measure Ly6C ratios at different ages. When possible, littermate controls were utilized for experiments. Both male and female mice and cells obtained from both genders were evaluated in this study. Mice were euthanized by CO_2_ inhalation. Mice were genotyped by standard PCR protocols.

All mice were maintained under specific pathogen-free conditions and all experiments were carried out in strict accordance with the recommendations outlined in the Guide for the Care and Use of Laboratory Animals from the National Institutes of Health. The protocol was approved by the Institutional Animal Care and Use Committee (IACUC) of the University of Maryland Baltimore School of Medicine (Protocol Number: 0218015).

## METHOD DETAILS

### FLOW CYTOMETRY (FACS ANALYSIS) AND CELL SORTING

Mice were sacrificed and whole blood samples were collected via cardiac puncture. After perfusion, spleen was macerated in Hank’s balanced salt solution (HBSS) and filtered through a 70-um cell strainer to obtain single cell suspension. Bone marrow cells were harvested by flushing femurs and tibias. Following hypotonic lysis with ACK buffer, single cell suspensions were pre-incubated with Mouse BD Fc Block™ (clone 2.4G2), then stained with a cocktail of fluorescently labeled antibodies (Abs) (see key resources table above) for 30 minutes at 4**°**C. All antibodies were titrated for optimal dilution. After staining, the cells were washed with 1x PBS and then viability dye 7-AAD was used to exclude dead cells from analysis. Flow cytometry was performed on a 3 laser (488 nm, 407 nm, 640 nm) Cytek Aurora spectral cytometer (Cytek Biosciences), and the spectral data were unmixed based on single color beads controls using SpectroFlo software, then analyzed using FCS Express 6 software (De Novo). ‘‘Fluorescence minus one’’ controls were used when necessary to confirm the correct gates.

Flow cytometry was performed according to ImmGen SOP (http://www.immgen.org/ATAC.Sort2017.pdf) (http://www.immgen.org/) and other previously published studies (Cossarizza et al., 2019; Mildner et al., 2016; Mildner et al., 2017; Yona et al., 2013), using the antibodies and gates indicated in Key Resources Table and Figure S2. Murine cell populations were defined as follows: **bone marrow cMOP:** Lin^-^ (Ter119^-^CD3^-^CD19^-^B220^-^ Ly6G^-^) CD115^+^CD117^+^CD135^-^; **bone marrow monocytes:** Lin^-^ (Ter119^-^CD3^-^CD19^-^B220^-^ Ly6G^-^) CD115^+^CD117^-^CD135^-^; **bone marrow Ly6C**^**hi**^ **monocytes:** Lin^-^ (Ter119^-^CD3^-^CD19^-^ B220^-^Ly6G^-^) CD115^+^CD117^-^CD135^-^CD11b^+^Ly6C^hi^; **bone marrow Ly6C**^**int**^ **monocytes:** Lin^-^ (Ter119^-^CD3^-^CD19^-^B220^-^Ly6G^-^) CD115^+^CD117^-^CD135^-^CD11b^+^Ly6C^int^; **bone marrow Ly6C**^**neg**^ **monocytes:** Lin^-^ (Ter119^-^CD3^-^CD19^-^B220^-^Ly6G^-^) CD115^+^CD117^-^CD135^-^ CD11b^+^Ly6C^neg^; **blood monocytes:** Lin^-^ (CD3^-^CD19^-^B220^-^Ly6G^-^) CD11b^+^CD115^+^; **blood Ly6C**^**hi**^ **monocytes:** Lin^-^ (CD3^-^CD19^-^B220^-^Ly6G^-^) CD115^+^CD11b^+^Ly6C^hi^; **blood Ly6C**^**int**^ **monocytes:** Lin^-^ (CD3^-^CD19^-^B220^-^Ly6G^-^) CD115^+^CD11b^+^Ly6C^int^; **blood Ly6C**^**neg**^ **monocytes:** Lin^-^ (CD3^-^CD19^-^B220^-^Ly6G^-^) CD115^+^CD11b^+^Ly6C^neg^; **spleen monocytes:** Lin^-^ (CD3^-^CD19^-^ B220^-^Ly6G^-^) CD45^+^CD11b^+^CD115^+^; **spleen Ly6C**^**hi**^ **monocytes**: Lin^-^ (CD3^-^CD19^-^B220^-^Ly6G^-^) CD45^+^CD115^+^CD11b^+^Ly6C^hi^; **spleen Ly6C**^**int**^ **monocytes**: Lin^-^ (CD3^-^CD19^-^B220^-^Ly6G^-^) CD45^+^CD115^+^CD11b^+^Ly6C^int^; **spleen Ly6C**^**neg**^ **monocytes**: Lin^-^ (CD3^-^CD19^-^B220^-^Ly6G^-^) CD45^+^CD115^+^CD11b^+^Ly6C^neg^; **bone marrow neutrophil**: CD11b^+^Ly6G^+^; **blood and spleen neutrophil**: CD3^-^CD19^-^CD11b^+^Ly6G^+^; **bone marrow, blood, and spleen B cells:** CD3^-^CD19^+^; **bone marrow, blood, and spleen T cell:** CD3^+^; **blood and spleen cDC:** CD45^+^Lin^-^ (CD3^-^ CD19^-^) MHCII^+^CD11c^+^; **blood and spleen pDC:** Lin^-^ (CD3^-^CD19^-^) PDCA1^+^B220^+^.

For experiments involving subset monocytes and OCs (osteoclast assay and RNA-sequencing), primary BM cells harvested from femurs and tibias were used to isolate monocytes using a Monocyte Isolation Kit (Miltenyi). Enriched monocytes were further stained with fluorescent Abs and sorted on FACS Aria II cell sorter (BD Biosciences, 4 lasers, 488 nm, 405 nm, 635 nm, 552 nm). Purity of the sorted monocyte cells were confirmed to be ∼90% and above. Data was analyzed using FCS Express 6 software (De Novo).

### MICRO-CT ANALYSIS

Femurs were scanned in a μCT 50 (Scanco Medical) at 70 kVp, 76 μA, 0.5 Al filter, with 1200 ms integration and 6 μm voxel dimension. Reconstructed images were analyzed in AnalyzePro 1.0 (AnalyzeDirect). Scans were calibrated to a standard curve of five known hydroxyapatite (HA) densities (mg/cm^3^). Calibrated images were reoriented and analyzed by standard cortical and trabecular bone algorithms (Bouxsein et al., 2010; Thumbigere-Math et al., 2018; Thumbigere-Math et al., 2019).

### HISTOLOGY

For histology, bones were fixed overnight in Bouin’s solution and demineralized in AFS (acetic acid, formaldehyde, sodium chloride) for 3-4 weeks and later embedded in paraffin, as previously described (Foster, 2012). Tartrate resistant acid phosphatase (TRAP) staining was performed on decalcified and deparaffinized tissues according to manufacturer’s instructions (Wako Chemical, Japan) (Thumbigere-Math et al., 2019). Number of TRAP+ osteoclasts along the bone perimeter of the epiphyseal growth plate was counted.

### CTX ASSAY

Blood was collected from mice and clotted for 2 hours at room temperature before centrifuge for 30 mins at 2000g. Later, serum was harvested and stored at –80°C for subsequent analysis. Serum C-terminal fragments of CTX were quantified using the RatLaps EIA assay (Immunodiagnostics Systems) according to the manufacturer’s instructions.

### IN VITRO OSTEOCLAST ASSAY

Osteoclast assays were performed as described previously (Thumbigere-Math et al., 2019). For experiments in Figures 2 and S3, briefly, primary BM cells were harvested from femurs and tibias of 8-12-week-old animals. After lysis of red blood cells using RBC lysis buffer (150 mM NH_4_Cl, 10 mM KHCO_3_, 0.1 mM EDTA, pH7.4), BM cells were cultured overnight in 100 mm tissue culture dishes (Corning) in osteoclast media (phenol red-free alpha-MEM (Gibco)) with 10% heat-inactivated fetal bovine serum (Hyclone), 25 units/mL penicillin/streptomycin (Invitrogen), 400 mM L-Glutamine (Invitrogen), and supplemented with 20 ng/ml M-CSF. The non-adherent cell population was recovered and re-plated in 12-well plates (Corning) at 5×10^5^ cells/cm^2^ in osteoclast media supplemented with 20 ng/ml M-CSF. The following day, cells were treated with 40 ng/mL RANKL (R&D Systems). Subsequently, media were replaced every 2 two days with 20 ng/ml M-CSF and 40 ng/mL RANKL to generate osteoclasts. Osteoclast formation was completed by 6 days after RANKL treatment.

For BMMs and OCs subset experiments in Figures 3-7, S4-7, FACS-sorted monocyte subsets were directly plated in 24-well plates at 2×10^5^ cells/cm^2^ or 48-well plates at 1×10^5^ cells/cm^2^ or 96-well plates at 0.3×10^5^ cells/cm^2^ and supplemented with 20 ng/ml M-CSF for 2 days, followed by treatment with 40 ng/mL RANKL. Subsequently, media were replaced every 2 two days with 20 ng/ml M-CSF and 40 ng/mL RANKL to generate osteoclasts. Osteoclast formation was completed by 4 days after RANKL treatment.

For tartrate resistant acid phosphatase (TRAP) staining, osteoclasts were rinsed in PBS and fixed with 4% paraformaldehyde for 20 minutes. The cells were then stained using Naphthol AS-MX phosphate and Fast Violet LB salt according to the protocol provided in BD Biosciences Technical Bulletin #445. Multinucleated (≥3 nuclei), TRAP-positive osteoclasts were counted in triplicate wells. Cells were imaged and photographed with light microscopy and the measurements were analyzed using NIH ImageJ (version 1.51).

The resorption activity of osteoclasts was examined using Osteo Assay Surface Plates (Corning) and cells were cultured as described above. Osteoclasts were removed by 5 minutes treatment with 10% bleach and washed with dH_2_O. The plates were then allowed to air dry completely at room temperature for 3–5 hours. The demineralized area was photographed by light microscopy and then analyzed using NIH ImageJ.

### IMMUNOBLOTTING

The assay was performed as described previously (Thumbigere-Math et al., 2019). Briefly, cell protein lysates were harvested using a lysis and extraction kit (ThermoFisher Scientific) supplemented with Halt Protease & Phosphatase Inhibitor Cocktail (ThermoFisher Scientific). Cytoplasmic and nucleoplasmic protein lysates were harvested using a Thermo NE-PER nuclear and cytoplasmic extraction kit, as directed by the manufacturer. Protein concentration was determined using the Bradford assay. Equal amounts of protein (20-30 μg) were loaded on NuPAGE Bis-Tris 4%–12% gradient polyacrylamide gels in the MOPS or MES buffer system (ThermoFisher Scientific) and subsequently electrotransferred from mini gels to a solid nitrocellulose membrane using the iBlot 2 system (ThermoFisher Scientific). Immediately nitrocellulose membranes were saturated for at least 1 hour at room temperature in blocking buffer (LICOR Odyssey). The membranes were then incubated with appropriate primary antibody (0.1–1 μg/ml) in blocking buffer with 0.1% Tween-20 for overnight at 4 °C. On the next day, the blot was incubated with anti-rabbit or anti-mouse HRP**−**conjugated secondary antibodies (Abcam) for 1 hour at room temperature. Antibody binding was detected using the ECL system (GE Healthcare). Images were acquired on film (Kodak) and cropped in Adobe Photoshop.

### QUANTITATIVE REAL-TIME PCR

The assay was performed as described previously (Thumbigere-Math et al., 2018; Thumbigere-Math et al., 2019). Briefly, total RNA was extracted from cultured cells using Trizol (Invitrogen). Complementary DNA was generated from 1 μg of total RNA with Superscript II RT (Invitrogen) and random hexamer primers (Promega). PCR was carried out by QuantStudio-3 Real-Time PCR System (Applied Biosystems) according to the manufacturer’s protocol. Transcript levels were normalized by Hprt and fold changes were calculated by the C_t_ method. Primers used are summarized in Supplemental Table S1.

### RNA-SEQ

RNA-seq was performed as described previously (Thumbigere-Math et al., 2019). Briefly, total RNA was first extracted from the unsorted or FACS-sorted cells using the RNeasy mini kit (Qiagen) and RNA integrity was evaluated with the Bioanalyzer RNA pico kit TapeStation (Agilent Technologies). Two hundred nanograms of total RNA was subsequently processed to generate RNA-seq libraries using NEBNext Poly(A) mRNA Magnetic Isolation and Ultra II Directional RNA Library Prep kit for Illumina (NEB E7490, E7760) according to manufacturer’s protocol. Library size distribution and yield were evaluated using the Picogreen analysis. Libraries were pooled at equimolar ratios and sequenced for 50-nt single-end cycling with HiSeq 3000 (Illumina).

### RNA-SEQ ANALYSIS

Unsorted Cells: Data from independent runs of 50 bp single-end reads were mapped to the mouse genome (UCSC genome browser, mm9 assembly) using TopHat 2.1.0. Mapped reads were analyzed using Partek GS v7 software (https://www.partek.com/) and normalized to RPKM (reads per kilobase of exon per million mapped). Differentially expressed genes were generated by one-way ANOVA analysis using Partek. PCA plots were generated using Partek against unfiltered RPKM data. Heat Maps were generated by plotting relative expression of group means of RPKM values from at least 3 biological replicates. The relative expression was calculated by subtracting the mean of group means and dividing by the standard deviation of the group means for each gene. For Global Heat Maps, genes in which at least one group mean RPKM was > 1 and the CV of group means across all the conditions was > 0.3 were included.

Sorted Ly6C^hi^, Ly6C^int^, and Ly6C^−^ Cells: Raw sequencing reads generated for each sample were analyzed using the CAVERN analysis pipeline (Shetty AC, 2019). Read quality was assessed using the FastQC toolkit (Andrews, 2010) to ensure good quality reads for downstream analyses. Reads were first aligned to the reference mouse genome (mm10 assembly) using HISAT2, a splice-aware alignment software for mapping next-generation sequencing reads (Kim et al., 2015). Reads were aligned using default parameters to generate the alignment BAM files. Read alignments were assessed to compute gene expression counts for each gene using the HTSeq count tool (Anders et al., 2015) and the mouse reference annotation (GRCm38.96). The raw read counts were normalized for library size and utilized to assess differential gene expression between WT and *Irf8 cKO* groups using the R package ‘DESeq2’ (Anders and Huber, 2010). P-values were generated using a Wald test implemented in DESeq2 and then corrected for multiple hypothesis testing using the Benjamini-Hochberg correction method. Significant differentially expressed genes between conditions were determined using a false discovery rate (FDR) of 1% and a minimum fold-change of 2X. PCA plots, volcano plots, and heatmaps were generated using R statistical analysis software using normalized read count data.

Gene set enrichment analysis (GSEA) from the Broad Institute (http://www.broad.mit.edu/gsea) was used against ANOVA results using Fold Change as the pre-ranked metric. Gene set enrichment analysis (GSEA) (Mootha et al., 2003; Subramanian et al., 2005) was used to determine the level of enrichment of selected gene sets within differential expression obtained from ANOVA comparisons of cells from different genotypes (Partek GS). The ANOVA fold changes were used as the ranking metric and GSEA Pre-ranked option was used with a “classic” enrichment score as per Broad Institute recommendations (http://software.broadinstitute.org/cancer/software/gsea/wiki/index.php/FAQ). 2000 permutations were performed. A reported p-value (and q-value) of 0 represents less than 1/2000 permutations. The RANKL-Responsive Genes list (n=238 genes) was developed using Ingenuity Pathway Analysis (IPA) to find genes whose gene expression is regulated by RANKL. Normalized BigWig files were generated using deepTools bamCoverage (Ramirez et al., 2016) and visualized using Integrative Genome Viewer (Broad Institute).

### CHIP-SEQ

Chromatin immunoprecipitation was performed as previously described (Dell’Orso et al., 2016). Briefly, cells were crosslinked in 1% formaldehyde at room temperature for 10□min and lysed in Farnham buffer (5□mM PIPES pH 8.0; 85□mM KCl; 0.5% NP-40) and subsequently in RIPA buffer (1x PBS; 1% NP-40; 0.5% sodium deoxycholate; 0.1% SDS). Chromatin was fragmented by sonication at high intensity and cycles of 30” ON/30” OFF with the Vibra-Cell Ultrasonic Liquid Processors for 10-20 minutes. Sheared chromatin was immobilized with Dynabeads Protein G (ThermoFisher) and immunoprecipitated with Anti-H3K27ac, H3K4me1, H3K4me3, H3K27me3, and PU.1 antibody and washed successively in buffer I (20□mM Tris HCl pH 8.0, 150□mM NaCl, 2□mM EDTA, 0.1% SDS, 1% Triton X-100); buffer II (20□mM Tris HCl pH 8.0, 500□mM NaCl, 2□mM EDTA, 0.1% SDS, 1% Triton X-100); buffer III (100□mM Tris HCl pH 8.0, 500□mM LiCl, 1% NP-40; 1% sodium deoxycholate). All washes were performed at 4°C for 3 to 5 min. Finally, aliquots of genomic DNA (input) and immunoprecipitated samples were treated with proteinase K, heated to induce de-cross linking, and purified using QIAquick PCR Purification Kit. After recovering purified DNA, 10ng or more of DNA was used to generate libraries according to the NEBNext RNA Library Prep Kit (New England Biolabs). Following the amplification step, DNA concentration was measured using the Picogreen analysis, and equimolar ratios of barcoded ChIPed DNA from each sample were pooled together. After barcoding, pooled DNA was sequenced (HiSeq 3000 Illumina) to achieve a minimum of 10^7^ aligned reads per sample.

### CHIP-SEQ ANALYSIS

Raw sequencing reads generated for each sample were aligned to the reference mouse genome (mm10 assembly) using Bowtie2 version 2.2.6 (Langmead and Salzberg, 2012). Reads were aligned with default parameters to generate the alignment BAM files and clonal reads were removed from further analysis. A minimum of 10 million uniquely mapped reads were obtained for each of the conditions. We used MACS2 (version 2.1.2) to identify peaks of ChIP-seq enrichment using default parameters. The appropriate input-DNA background was set with the ‘--control’ parameter. In all cases, redundant reads were removed, and only one mapped read to each unique regions of the genome was kept and used in peak calling. ChIP-seq data were visualized by preparing custom tracks for the UCSC Genome browser.

Read alignments were used to identify enrichments of histone marks (H3K4me1, H3K4me3, H3K27ac, H3K27me3) and transcription factors (PU.1 and IRF8) within the 4kb flanking region of the TSS for all the differentially expressed genes identified from the RNA-Seq experiments (sorted Ly6C^hi^, Ly6C^int^, and Ly6C^−^ cells). Previously published ChIP-seq profile immunoprecipitated from murine BMMs (Mancino et al., 2015) was used to map genome-wide IRF8 binding sites. Downstream analyses were done using custom R programing to assess the relationship between gene expression and each of the histone marks. Chromatin state analysis integrated the fold-enrichments from the gene expression with the fold-enrichment from each of the histone marks to identify genes in active promoter state, strong enhancer state, or repressor state. Active promoters were identified according to proximity to transcription start sites (TSS) (<2 kb) and H3K4me3, while active enhancers were defined by their distance from TSS (>2 kb) and enrichment of both H3K4me1 and H3K27ac (Creyghton et al., 2010; Heintzman et al., 2007). Transcriptional repressors were identified by their distance from TSS (<2 kb) and enrichment of H3K27me3. Profiles and HeatMap, as well as other downstream analyses, were done using R programing.

Motif Enrichment Analysis: de novo and known motif analysis were performed within the identified active promoter and active enhancer regions, using command findMotifsGenome.pl from HOMER package version 4.1.1 (Heinz et al., 2010). Peak sequences were compared to random genomic fragments of the same size and normalized G+C content to identify motifs enriched in the targeted sequences.

### QUANTIFICATION AND STATISTICAL ANALYSIS

All data are presented as mean±SD, unless stated otherwise. Statistical tests were selected based on appropriate assumptions with respect to data distribution and variance characteristics. Data were analyzed by one-way ANOVA and post-hoc Tukey’s multiple comparison test using GraphPad Prism 6 (GraphPad Software, Inc.). Where appropriate (comparison of two groups only), two-tailed paired or unpaired Student’s *t* tests were performed. Throughout the paper, error bars indicate mean ± STD and asterisks denote statistical significance (\**P* < 0.05; \*\**P* < 0.01; \*\*\**P* < 0.001; \*\*\*\**P* < 0.0001; NS, not significant). *n* represents the number of biological replicates. For *in vitro* and *in vivo* experiments, number of biological replicates in each group is specified in the figures. As for *in vitro* experiments, samples were excluded from analysis only in case of clear technical problem. Bone loss measurements and histological analysis were performed in a blinded fashion, whereas the conduct of experiments and assessment of other outcomes were not blinded to the study personnel. All *in vitro* experiments were performed three or more times (in triplicates) to ensure reproducibility of the observations. RNA-seq was performed once with 3 biological replicates, whereas ChIP-seq was performed once. The concordance between RNA-seq biological replicates was very high (R2 range, 0.988–0.998).

## Notes

### Competing Interest Statement

The authors have declared no competing interest.

