## Supplemental Figures for "Monocyte Subsets with High Osteoclastogenic Potential and Their Epigenetic Regulation Orchestrated by IRF8"

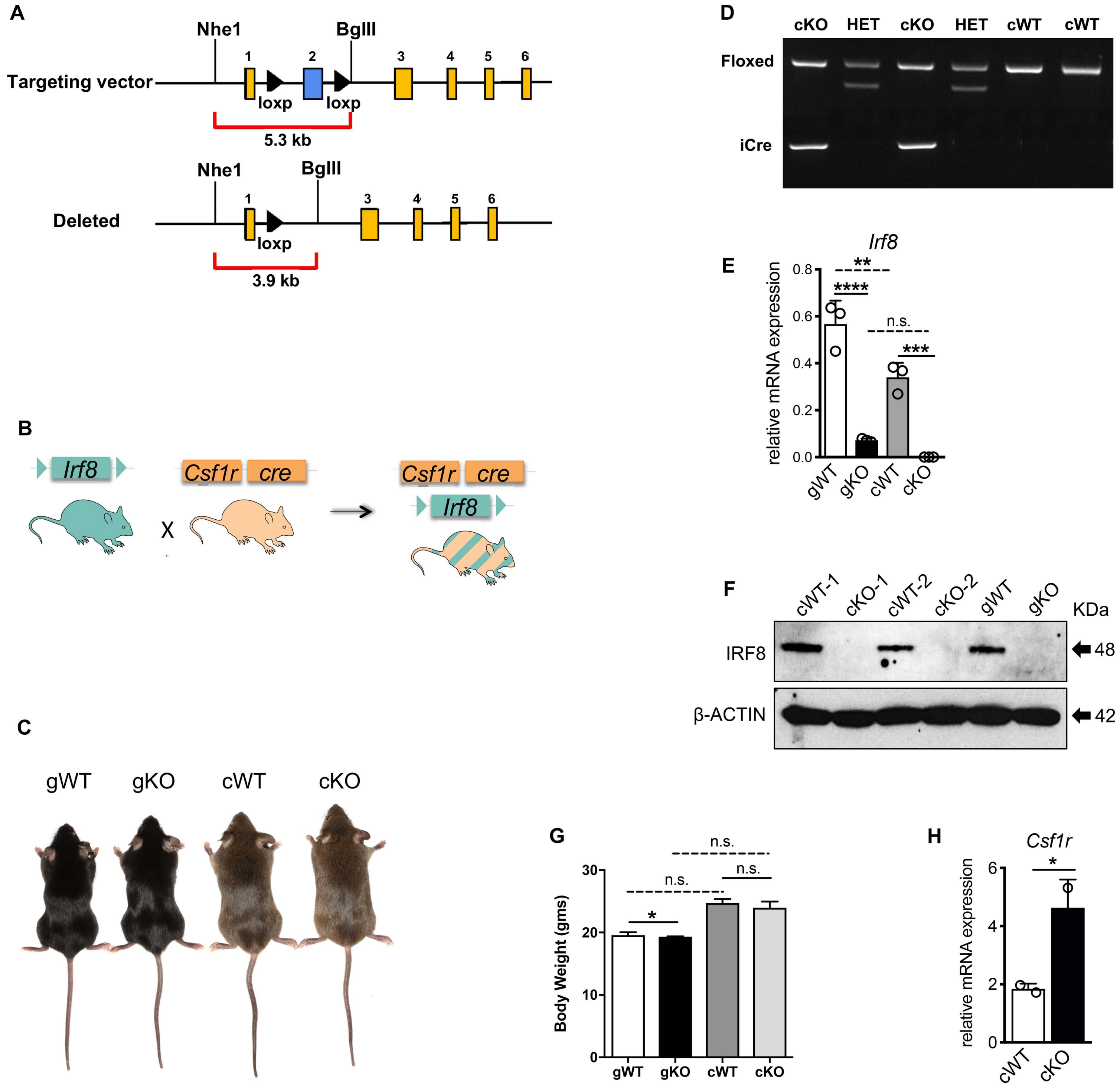

A
Nhe1
BglII
Targeting vector
loxp
loxp
5.3 kb
Deleted
3.9 kb
D
cKO HET cKO HET cWT cWT
Floxed
iCre
E
Irf8
relative mRNA expression
gWT gKO cWT cKO
n.s.
B
Irf8
Csf1r
cre
X
F
cWT-1 cKO-1 cWT-2 cKO-2 gWT gKO
KDa
IRF8
β-ACTIN
48
42
C
gWT gKO cWT cKO
G
Body Weight (gms)
n.s.
H
Csf1r
relative mRNA expression
cWT cKO

### **Figure S1. Generation of *Irf8 cKO* mice**

(A) Targeting strategy to conditionally delete exon 2 of *Irf8*.

(B) Schematic of the crosses to generate *Irf8 cKO* mice.

(C) Photographic images of *Irf8 gWT, gKO, cWT* and *cKO* mice. WT's and KO's are from respective 9-week-old littermates.

(D) PCR Genotyping of *Irf8 cKO* and *cWT* mice.

(E) qPCR analysis shows lack of IRF8 transcript in BMMs of Irf8 *cKO* mice, confirming Csf1r-cre mediated *Irf8* deletion. Other genotypes serve as control groups.

(F) Immunoblot shows lack of IRF8 protein in BMMs of Irf8 *cKO* mice, confirming Csf1r-cre mediated IRF8 deletion. Other genotypes serve as control groups.

(G) Body weight of 9-week-old *Irf8 gWT, gKO, cWT* and *cKO* mice (n = 5 mice per genotype).

(H) BMMs from *Irf8 cWT* and *cKO* mice were tested for mRNA expression of *Csf1r* by RT-qPCR analysis.

(I) Error bars indicate mean ± STD. *$p < 0.05$, **$p < 0.01$, ***$p < 0.001$, ****$p < 0.0001$, and n.s. = non-significant.

See also Table S1 for qPCR primer sequences.

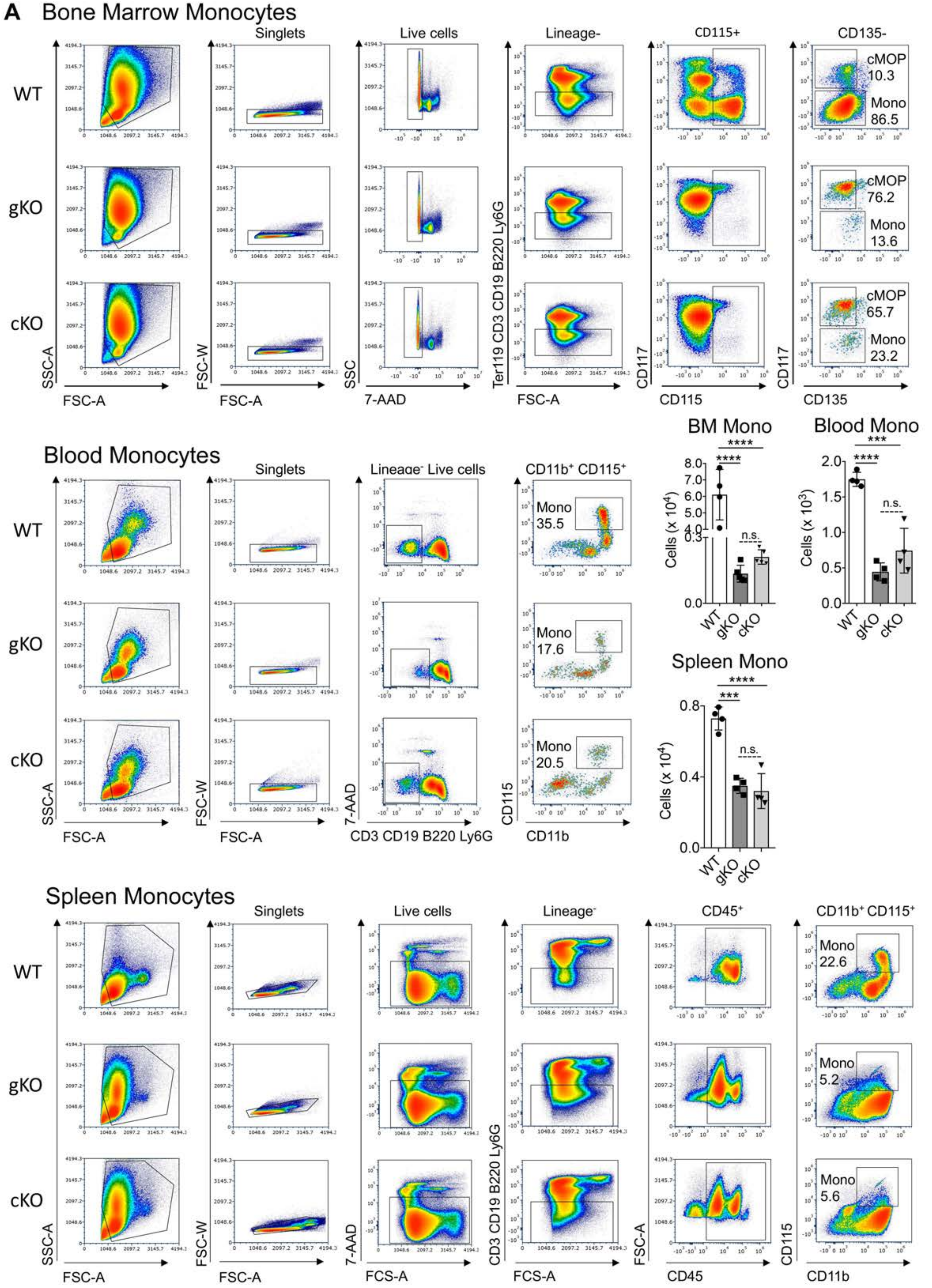
A
Bone Marrow Monocytes
Singlets
Live cells
Lineage-
CD115+
CD135-
WT
gKO
cKO
cMOP
10.3
Mono
86.5
cMOP
76.2
Mono
13.6
cMOP
65.7
Mono
23.2
SSC-A
FSC-A
FSC-W
FSC-A
SSC
7-AAD
Ter119 CD3 CD19 B220 Ly6G
FSC-A
CD117
CD115
CD117
CD135
BM Mono
Blood Mono
Blood Monocytes
Singlets
Lineage⁻ Live cells
CD11b⁺ CD115⁺
Mono
35.5
Mono
17.6
Mono
20.5
SSC-A
FSC-A
FSC-W
FSC-A
7-AAD
CD3 CD19 B220 Ly6G
CD115
CD11b
Cells (x 10⁴)
Cells (x 10³)
n.s.
Spleen Mono
Cells (x 10⁴)
Spleen Monocytes
Singlets
Live cells
Lineage⁻
CD45⁺
CD11b⁺ CD115⁺
Mono
22.6
Mono
5.2
Mono
5.6
SSC-A
FSC-A
FSC-W
FSC-A
7-AAD
FCS-A
CD3 CD19 B220 Ly6G
FCS-A
FSC-A
CD45
CD115
CD11b

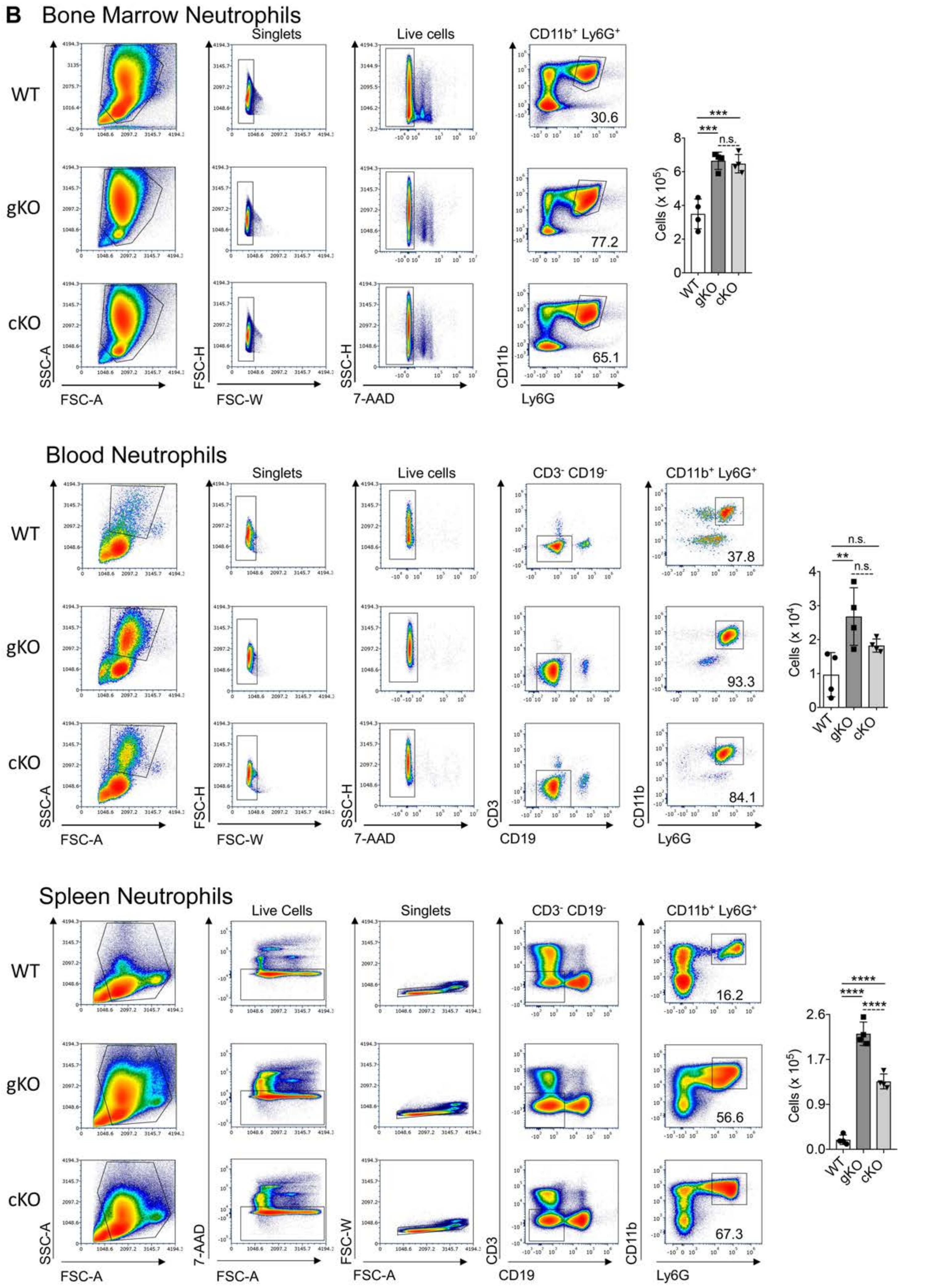
B
Bone Marrow Neutrophils
Singlets
Live cells
CD11b⁺ Ly6G⁺
WT
gKO
cKO
30.6
77.2
65.1
SSC-A
FSC-A
FSC-H
FSC-W
SSC-H
7-AAD
CD11b
Ly6G
***
***
n.s.
Cells (x 10^5)
Blood Neutrophils
Singlets
Live cells
CD3⁻ CD19⁻
CD11b⁺ Ly6G⁺
37.8
93.3
84.1
CD3
CD19
n.s.
**
n.s.
Cells (x 10^4)
Spleen Neutrophils
Live Cells
Singlets
CD3⁻ CD19⁻
CD11b⁺ Ly6G⁺
16.2
56.6
67.3
****
****
****
Cells (x 10^5)

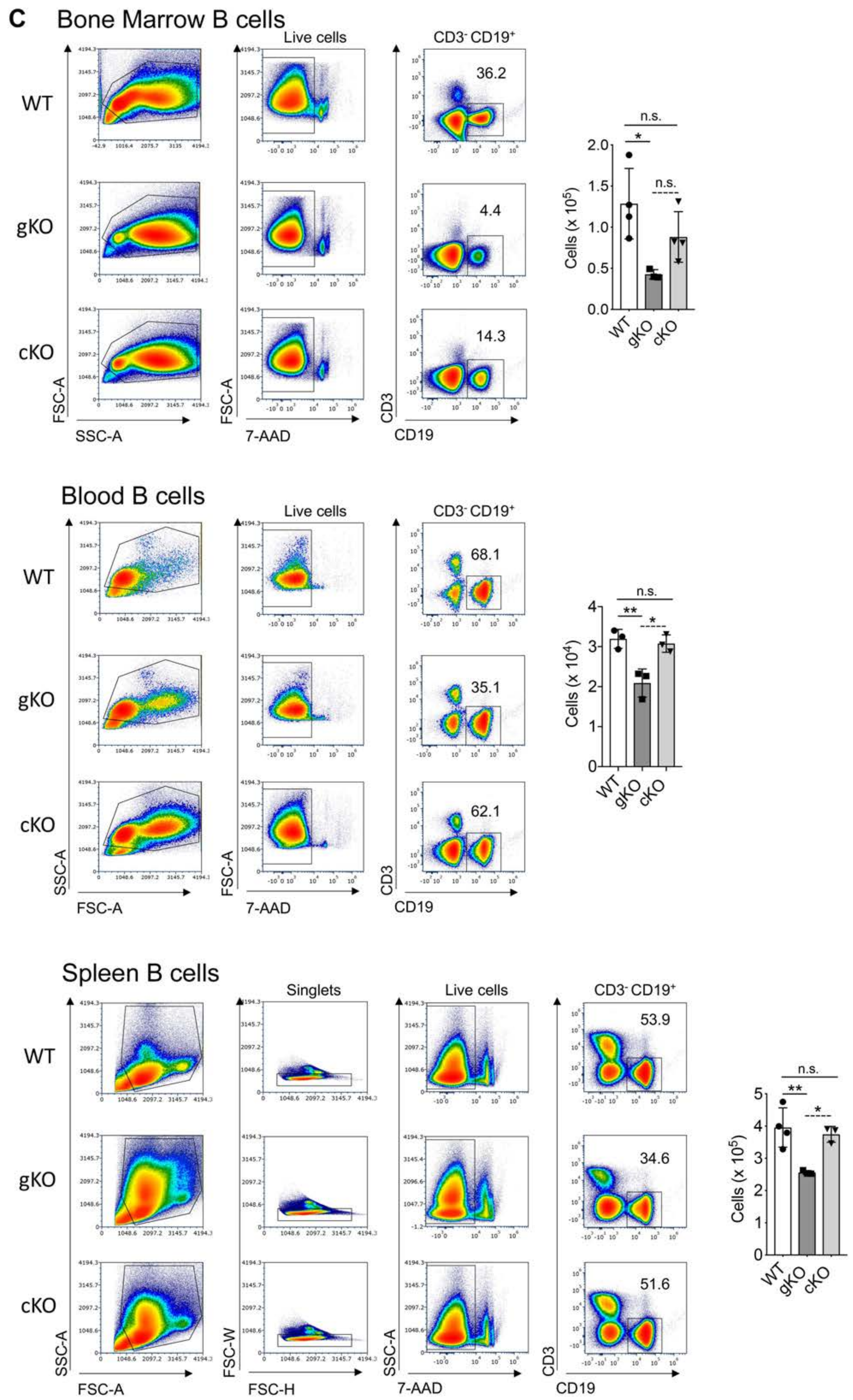
C
Bone Marrow B cells
Live cells
CD3- CD19+
WT
gKO
cKO
36.2
4.4
14.3
FSC-A
SSC-A
7-AAD
CD3
CD19
n.s.
*
Cells (x 10^5)
Blood B cells
68.1
35.1
62.1
**
Cells (x 10^4)
Spleen B cells
Singlets
53.9
34.6
51.6
FSC-W
FSC-H
Cells (x 10^5)

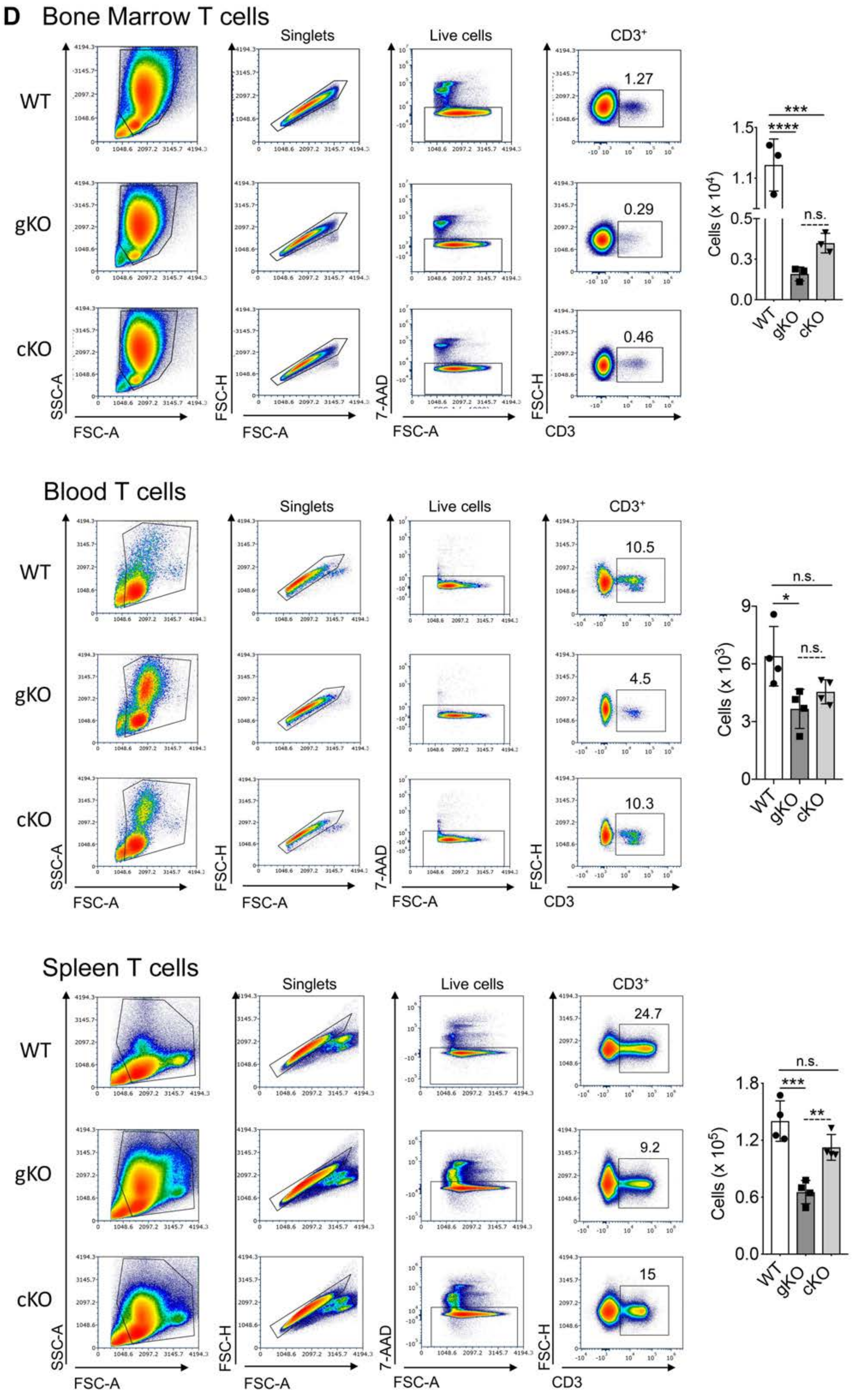

D
Bone Marrow T cells
Singlets
Live cells
CD3+
WT
gKO
cKO
1.27
0.29
0.46
SSC-A
FSC-A
FSC-H
7-AAD
CD3
Cells (x 10^4)
Blood T cells
10.5
4.5
10.3
Cells (x 10^3)
n.s.
Spleen T cells
24.7
9.2
15
Cells (x 10^5)

**E**

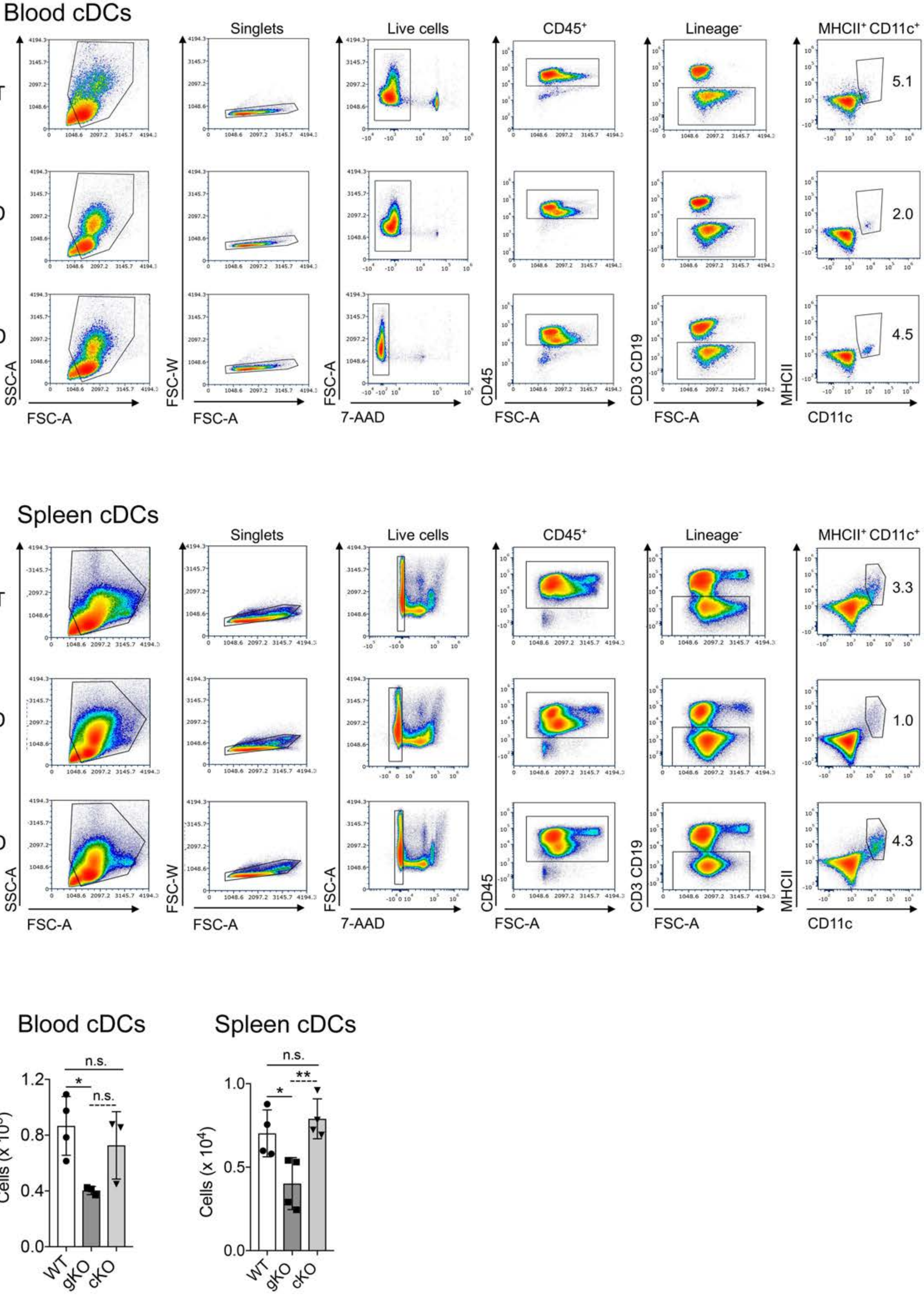
Blood cDCs
Singlets
Live cells
CD45+
Lineage-
MHCII+ CD11c+
WT
gKO
cKO
5.1
2.0
4.5
SSC-A
FSC-A
FSC-W
7-AAD
CD45
CD3 CD19
MHCII
CD11c
Spleen cDCs
3.3
1.0
4.3
Blood cDCs
Spleen cDCs
n.s.
*
**
Cells (x 10^3)
Cells (x 10^4)

F

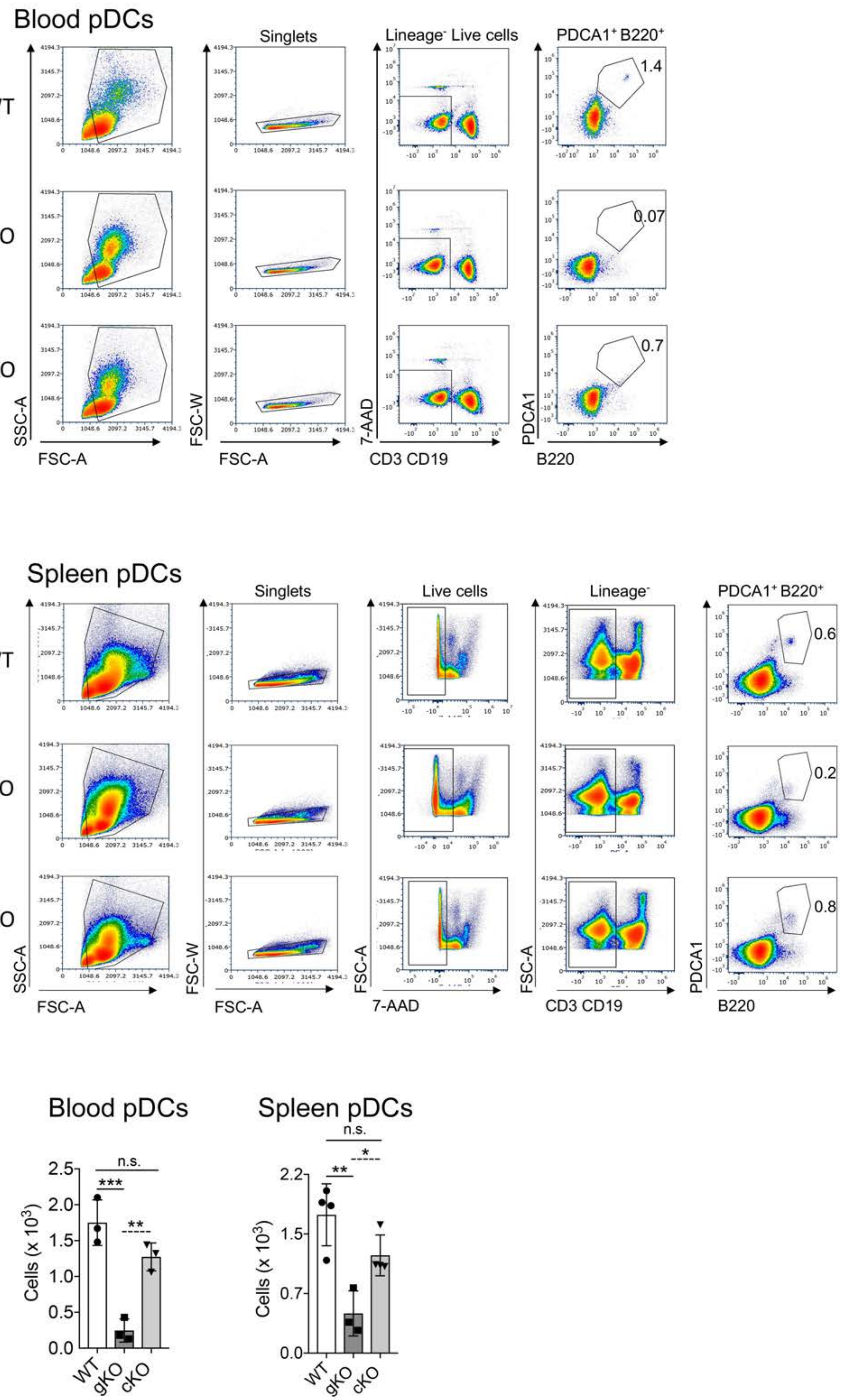

Blood pDCs
Singlets
Lineage⁻ Live cells
PDCA1⁺ B220⁺
WT
gKO
cKO
1.4
0.07
0.7
SSC-A
FSC-A
FSC-W
FSC-A
7-AAD
CD3 CD19
PDCA1
B220
Spleen pDCs
Singlets
Live cells
Lineage⁻
PDCA1⁺ B220⁺
WT
gKO
cKO
0.6
0.2
0.8
SSC-A
FSC-A
FSC-W
FSC-A
FSC-A
7-AAD
FSC-A
CD3 CD19
PDCA1
B220
Blood pDCs
n.s.
***
**
Cells (x 10³)
WT
gKO
cKO
Spleen pDCs
n.s.
*
**
Cells (x 10³)
WT
gKO
cKO

**Figure S2, related to Figure 1. Comprehensive flow cytometric analysis of HSCs in *Irf8 cKO* Mice**

Gating strategy used for identification of (A) Monocytes, (B) Neutrophils, (C) B cells, (D) T cells, (E) Conventional dendritic cells (cDCs), and (F) Plasmacytoid dendritic cells (pDCs) in bone marrow, peripheral blood, and spleen samples of *Irf8 WT, gKO* and *cKO* mice. Pseudocolor plots show cell population in percentages and bar graphs show absolute counts. Figures are representative of 4-5 independent experiments (n = 4-5 mice per genotype).

Error bars indicate mean ± STD. *$p < 0.05$, **$p < 0.01$, ***$p < 0.001$, ****$p < 0.0001$, and n.s. = non-significant.

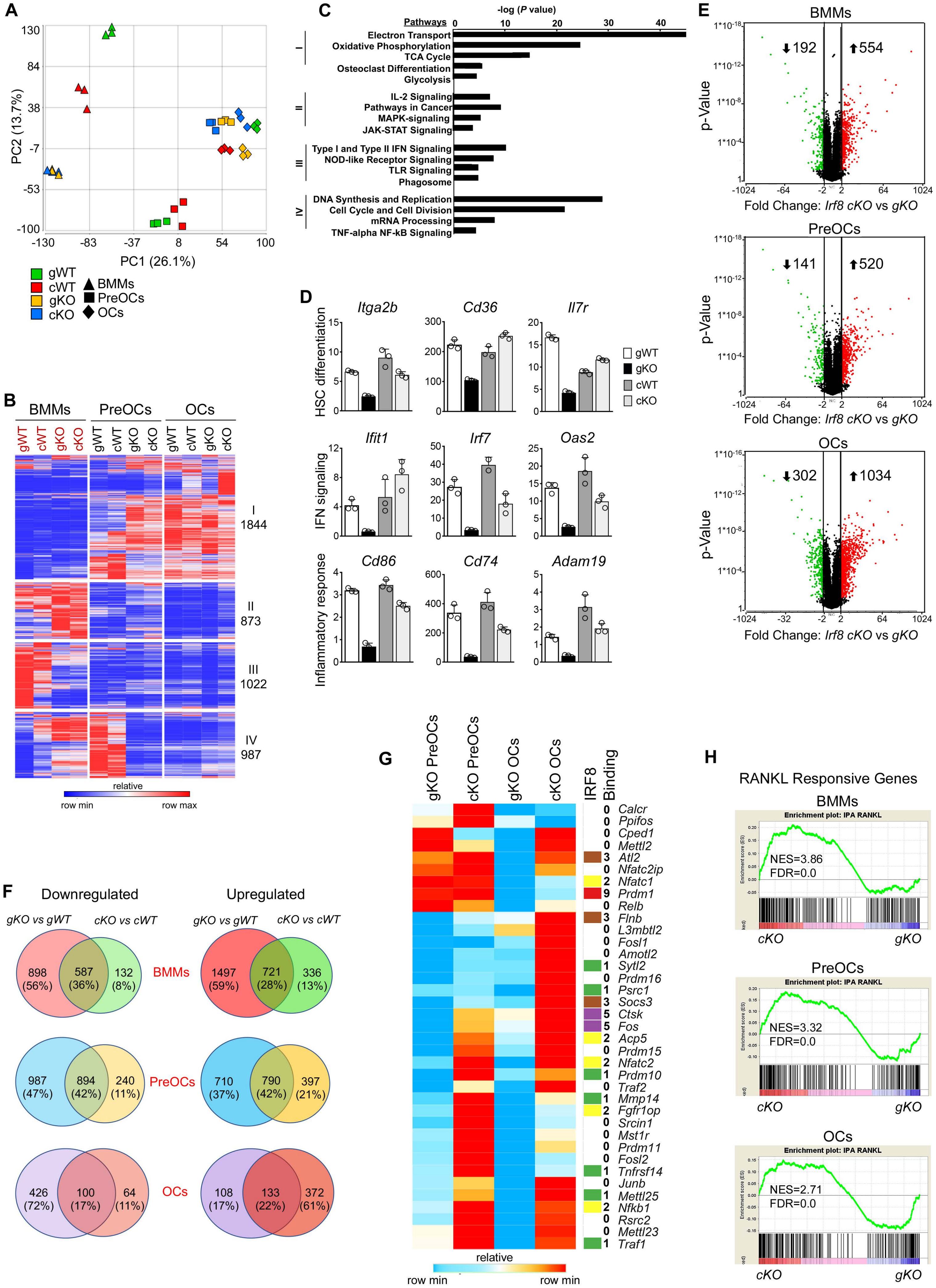
A
PC2 (13.7%)
PC1 (26.1%)
130
84
38
-7
-53
-100
-130
-83
-37
8
54
100
gWT
cWT
gKO
cKO
BMMs
PreOCs
OCs
B
BMMs
PreOCs
OCs
gWT
cWT
gKO
cKO
I
1844
II
873
III
1022
IV
987
relative
row min
row max
C
-log (P value)
Pathways
0
10
20
30
40
Electron Transport
Oxidative Phosphorylation
TCA Cycle
Osteoclast Differentiation
Glycolysis
IL-2 Signaling
Pathways in Cancer
MAPK-signaling
JAK-STAT Signaling
Type I and Type II IFN Signaling
NOD-like Receptor Signaling
TLR Signaling
Phagosome
DNA Synthesis and Replication
Cell Cycle and Cell Division
mRNA Processing
TNF-alpha NF-kB Signaling
D
HSC differentiation
Itga2b
Cd36
Il7r
IFN signaling
Ifit1
Irf7
Oas2
Inflammatory response
Cd86
Cd74
Adam19
gWT
gKO
cWT
cKO
E
BMMs
PreOCs
OCs
p-Value
Fold Change: Irf8 cKO vs gKO
192
554
141
520
302
1034
F
Downregulated
Upregulated
gKO vs gWT
cKO vs cWT
BMMs
PreOCs
OCs
898 (56%)
587 (36%)
132 (8%)
1497 (59%)
721 (28%)
336 (13%)
987 (47%)
894 (42%)
240 (11%)
710 (37%)
790 (42%)
397 (21%)
426 (72%)
100 (17%)
64 (11%)
108 (17%)
133 (22%)
372 (61%)
G
gKO PreOCs
cKO PreOCs
gKO OCs
cKO OCs
IRF8 Binding
0 Calcr
0 Ppifos
0 Cped1
0 Mettl2
3 Atl2
0 Nfatc2ip
2 Nfatc1
9 Prdm1
0 Relb
3 Flnb
0 L3mbtl2
0 Fosl1
0 Amotl2
1 Sytl2
0 Prdm16
1 Psrc1
3 Socs3
5 Ctsk
5 Fos
2 Acp5
0 Prdm15
2 Nfatc2
1 Prdm10
0 Traf2
1 Mmp14
2 Fgfr1op
0 Srcin1
0 Mst1r
0 Prdm11
0 Fosl2
1 Tnfrsf14
0 Junb
1 Mettl25
2 Nfkb1
0 Rsrc2
0 Mettl23
1 Traf1
relative
row min
row min
H
RANKL Responsive Genes
BMMs
PreOCs
OCs
Enrichment plot: IPA RANKL
NES=3.86
FDR=0.0
NES=3.32
FDR=0.0
NES=2.71
FDR=0.0
cKO
gKO

**Figure S3, related to Figure 2. *Irf8 cKO* Mice Provide Novel Osteoclast Transcriptomic Results**

(A) PCA of RNA-seq data from *Irf8 gWT, gKO, cWT* and *cKO* BMMs and RANKL treated Pre-osteoclasts (PreOCs) and mature osteoclasts (OCs).

(B) Hierarchical clustered heat map of transcripts regulated in *Irf8 gWT, gKO, cWT* and *cKO* cells during osteoclast differentiation process. Differential genes clustered into four major groups.

(C) Enriched functions for each cluster.

(D) Expression levels from individual genes identified by the cluster analysis depicted in (B). Shown are the mean sequence reads ± STD.

(E) Volcano plot illustrates significant differences in gene expression pattern between *Irf8 cKO* vs *gKO* BMMs, PreOCs and OCs (one-way ANOVA, >2-fold change, FDR<0.01). Analysis includes direct comparison between *Irf8 cKO* and *gKO* transcriptome. Red dots=upregulated genes, green dots=downregulated genes, and black dots=no significant change.

(F) Venn diagram depicting overlaps between *Irf8 gKO* vs *gWT* (fold change >2 or < -2, FDR<0.01) and *Irf8 cKO* vs *cWT* (fold change >2 or < -2, FDR<0.01) BMMs, PreOCs and OCs.

(G) Expression level changes for selected osteoclast-specific markers (n=37) in *Irf8 cKO* and *gKO* PreOCs and OCs. Shaded heat map on the right indicates the presence of one or more IRF8 binding sites.

(H) GSEA analysis shows strong enrichment of osteoclast-specific signatures in *Irf8 cKO* vs *gKO* BMMs, PreOCs and OCs.

(I) All data includes results from 3 biological replicates for each genotype. Data in heat maps are presented as standardized RPKM scale.

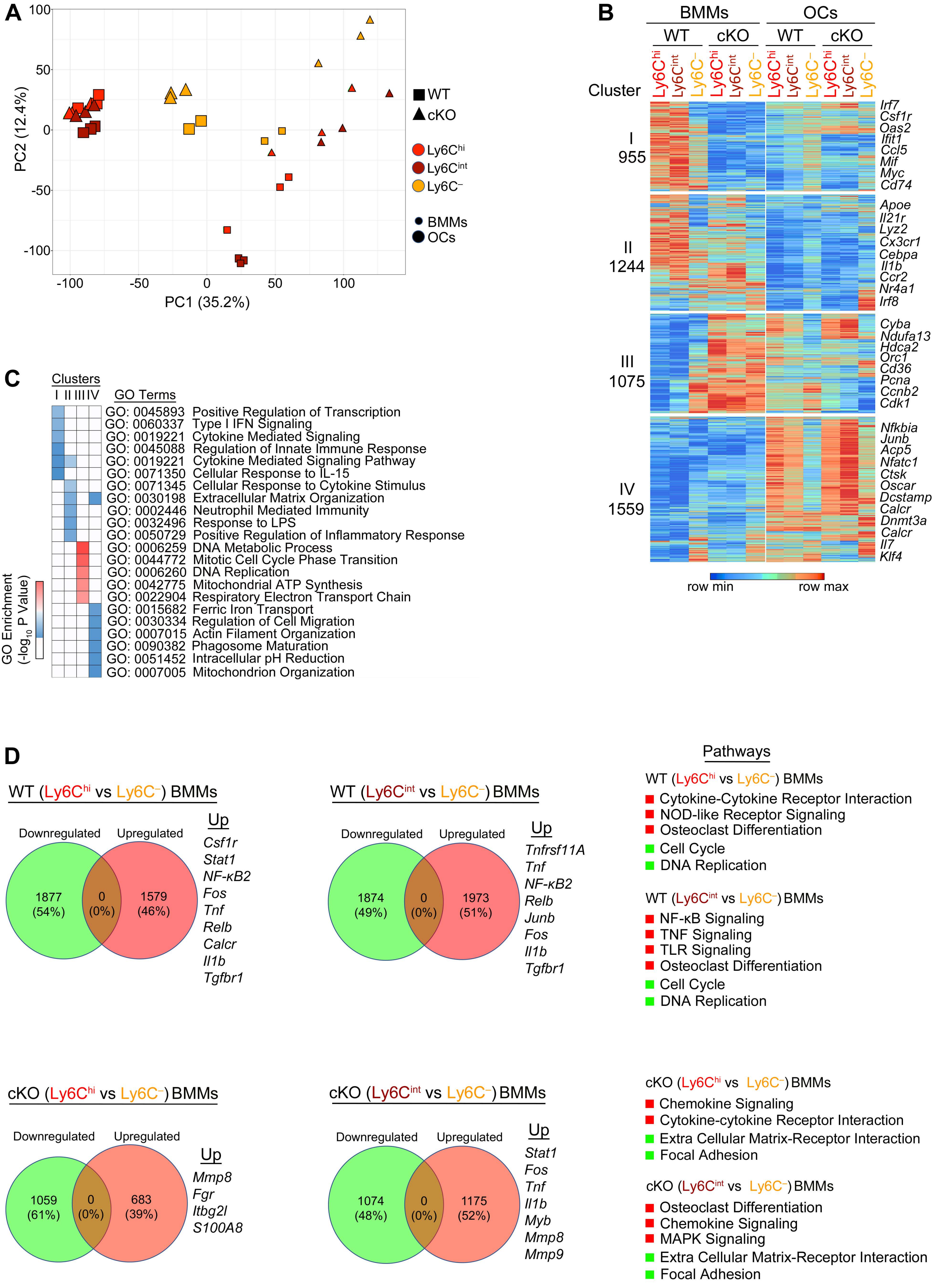
A
PC2 (12.4%)
PC1 (35.2%)
WT
cKO
Ly6Chi
Ly6Cint
Ly6C–
BMMs
OCs
B
BMMs
OCs
WT
cKO
WT
cKO
Cluster
Ly6Chi Ly6Cint Ly6C– Ly6Chi Ly6Cint Ly6C– Ly6Chi Ly6Cint Ly6C– Ly6Chi Ly6Cint Ly6C–
I
955
II
1244
III
1075
IV
1559
Irf7
Csf1r
Oas2
Ifit1
Ccl5
Mif
Myc
Cd74
Apoe
Il21r
Lyz2
Cx3cr1
Cebpa
Il1b
Ccr2
Nr4a1
Irf8
Cyba
Ndufa13
Hdca2
Orc1
Cd36
Pcna
Ccnb2
Cdk1
Nfkbia
Junb
Acp5
Nfatc1
Ctsk
Oscar
Dcstamp
Calcr
Dnmt3a
Calcr
Il7
Klf4
row min
row max
C
Clusters
I II III IV
GO Terms
GO: 0045893 Positive Regulation of Transcription
GO: 0060337 Type I IFN Signaling
GO: 0019221 Cytokine Mediated Signaling
GO: 0045088 Regulation of Innate Immune Response
GO: 0019221 Cytokine Mediated Signaling Pathway
GO: 0071350 Cellular Response to IL-15
GO: 0071345 Cellular Response to Cytokine Stimulus
GO: 0030198 Extracellular Matrix Organization
GO: 0002446 Neutrophil Mediated Immunity
GO: 0032496 Response to LPS
GO: 0050729 Positive Regulation of Inflammatory Response
GO: 0006259 DNA Metabolic Process
GO: 0044772 Mitotic Cell Cycle Phase Transition
GO: 0006260 DNA Replication
GO: 0042775 Mitochondrial ATP Synthesis
GO: 0022904 Respiratory Electron Transport Chain
GO: 0015682 Ferric Iron Transport
GO: 0030334 Regulation of Cell Migration
GO: 0007015 Actin Filament Organization
GO: 0090382 Phagosome Maturation
GO: 0051452 Intracellular pH Reduction
GO: 0007005 Mitochondrion Organization
GO Enrichment (-log10 P Value)
D
WT (Ly6Chi vs Ly6C–) BMMs
Downregulated
Upregulated
1877 (54%)
0 (0%)
1579 (46%)
Up
Csf1r
Stat1
NF-κB2
Fos
Tnf
Relb
Calcr
Il1b
Tgfbr1
WT (Ly6Cint vs Ly6C–) BMMs
Downregulated
Upregulated
1874 (49%)
0 (0%)
1973 (51%)
Up
Tnfrsf11A
Tnf
NF-κB2
Relb
Junb
Fos
Il1b
Tgfbr1
Pathways
WT (Ly6Chi vs Ly6C–) BMMs
Cytokine-Cytokine Receptor Interaction
NOD-like Receptor Signaling
Osteoclast Differentiation
Cell Cycle
DNA Replication
WT (Ly6Cint vs Ly6C–) BMMs
NF-κB Signaling
TNF Signaling
TLR Signaling
Osteoclast Differentiation
Cell Cycle
DNA Replication
cKO (Ly6Chi vs Ly6C–) BMMs
Downregulated
Upregulated
1059 (61%)
0 (0%)
683 (39%)
Up
Mmp8
Fgr
Itbg2l
S100A8
cKO (Ly6Cint vs Ly6C–) BMMs
Downregulated
Upregulated
1074 (48%)
0 (0%)
1175 (52%)
Up
Stat1
Fos
Tnf
Il1b
Myb
Mmp8
Mmp9
cKO (Ly6Chi vs Ly6C–) BMMs
Chemokine Signaling
Cytokine-cytokine Receptor Interaction
Extra Cellular Matrix-Receptor Interaction
Focal Adhesion
cKO (Ly6Cint vs Ly6C–) BMMs
Osteoclast Differentiation
Chemokine Signaling
MAPK Signaling
Extra Cellular Matrix-Receptor Interaction
Focal Adhesion

**Figure S4, related to Figure 5 and Figure S5. Transcriptional Profiling of Ly6C$^{hi}$, Ly6C$^{int}$, and Ly6C$^{–}$ Monocytes and Osteoclasts in WT and *Irf8 cKO* mice**

(A) Principal Component Analysis (PCA) of RNA-seq replicates from WT and *Irf8 cKO* Ly6C$^{hi}$, Ly6C$^{int}$, and Ly6C$^{–}$ subsets before (BMMs) and after 4 days of RANKL treatment (OCs).

(B) Hierarchical clustered heat map of 4,833 RANKL-responsive genes in Ly6C$^{hi}$, Ly6C$^{int}$, and Ly6C$^{–}$ BMMs and OCs in WT and *Irf8 cKO* mice. Differential genes clustered into four major groups. Data in heat maps are presented as standardized RPKM scale.

(C) Heatmap showing the *P*-value significance of GO term enrichment for genes in each cluster.

(D) Venn diagram shows transcriptomic changes between Ly6C$^{hi}$ vs Ly6C$^{–}$ BMMs and Ly6C$^{int}$ vs Ly6C$^{–}$ BMMs in WT and *Irf8 cKO* mice. Enriched OC-specific genes and enriched pathways (KEGG) in each group are presented as well.

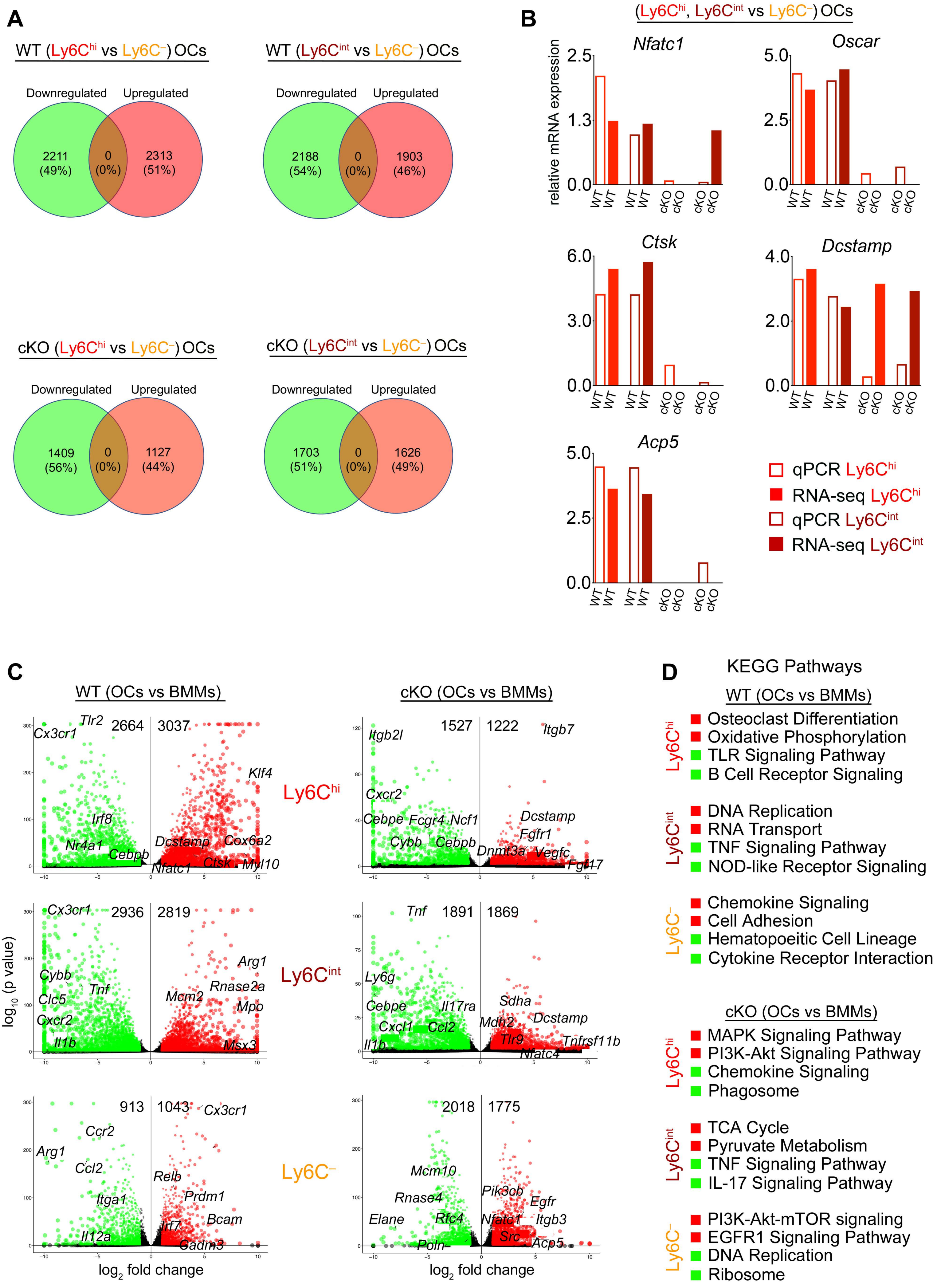
A
WT (Ly6Chi vs Ly6C−) OCs
Downregulated
Upregulated
2211 (49%)
0 (0%)
2313 (51%)
WT (Ly6Cint vs Ly6C−) OCs
2188 (54%)
0 (0%)
1903 (46%)
cKO (Ly6Chi vs Ly6C−) OCs
1409 (56%)
0 (0%)
1127 (44%)
cKO (Ly6Cint vs Ly6C−) OCs
1703 (51%)
0 (0%)
1626 (49%)
B
(Ly6Chi, Ly6Cint vs Ly6C−) OCs
relative mRNA expression
Nfatc1
Oscar
Ctsk
Dcstamp
Acp5
WT
cKO
qPCR Ly6Chi
RNA-seq Ly6Chi
qPCR Ly6Cint
RNA-seq Ly6Cint
C
WT (OCs vs BMMs)
cKO (OCs vs BMMs)
Ly6Chi
Ly6Cint
Ly6C−
2664
3037
1527
1222
2936
2819
1891
1869
913
1043
2018
1775
log10 (p value)
log2 fold change
Cx3cr1
Tlr2
Klf4
Irf8
Nr4a1
Cebpb
Dcstamp
Cox6a2
Nfatc1
Ctsk
Myl10
Itgb2l
Itgb7
Cxcr2
Cebpe
Fcgr4
Ncf1
Cybb
Fgfr1
Dnmt3a
Vegfc
Fgf17
Arg1
Tnf
Clc5
Rnase2a
Mcm2
Mpo
Il1b
Msx3
Ly6g
Il17ra
Sdha
Cxcl1
Ccl2
Mdh2
Tlr9
Tnfrsf11b
Nfatc4
Ccr2
Relb
Prdm1
Itga1
Bcam
Irf7
Il12a
Cadm3
Mcm10
Pik3cb
Egfr
Rnase4
Elane
Rfc4
Itgb3
Src
Acp5
Poln
D
KEGG Pathways
WT (OCs vs BMMs)
Ly6Chi
Osteoclast Differentiation
Oxidative Phosphorylation
TLR Signaling Pathway
B Cell Receptor Signaling
Ly6Cint
DNA Replication
RNA Transport
TNF Signaling Pathway
NOD-like Receptor Signaling
Ly6C−
Chemokine Signaling
Cell Adhesion
Hematopoeitic Cell Lineage
Cytokine Receptor Interaction
cKO (OCs vs BMMs)
Ly6Chi
MAPK Signaling Pathway
PI3K-Akt Signaling Pathway
Chemokine Signaling
Phagosome
Ly6Cint
TCA Cycle
Pyruvate Metabolism
TNF Signaling Pathway
IL-17 Signaling Pathway
Ly6C−
PI3K-Akt-mTOR signaling
EGFR1 Signaling Pathway
DNA Replication
Ribosome

**Figure S5, related to Figure 5 and Figure S4. Transcriptomic Analysis of Ly6C$^{hi}$, Ly6C$^{int}$, and Ly6C$^{–}$ Monocytes and Osteoclasts**

(A) Venn diagram shows transcriptomic changes between Ly6C$^{hi}$ vs Ly6C$^{–}$ OCs and Ly6C$^{int}$ vs Ly6C$^{–}$ OCs in WT and *Irf8 cKO* mice.

(B) Enriched OC-specific genes in Ly6C$^{hi}$ vs Ly6C$^{–}$ OCs and Ly6C$^{int}$ vs Ly6C$^{–}$ OCs in WT and *Irf8 cKO* mice. Both RNA-Seq and RT-qPCR analyses show similar expression changes. Results indicate that OC-specific genes (except *Dcstamp* RNA-seq) were almost equally expressed in all three subset OCs of *Irf8 cKO* mice.

(C) Volcano plot of transcriptomic changes between OCs vs BMMs in WT and *Irf8* mice. Color dots (red=up, green=down, back=no significant change) correspond to genes with >2-fold change, FDR<0.01 (one-way ANOVA Analysis).

(D) Enriched KEGG pathways in each group.

See also Table S1 for qPCR primer sequences.

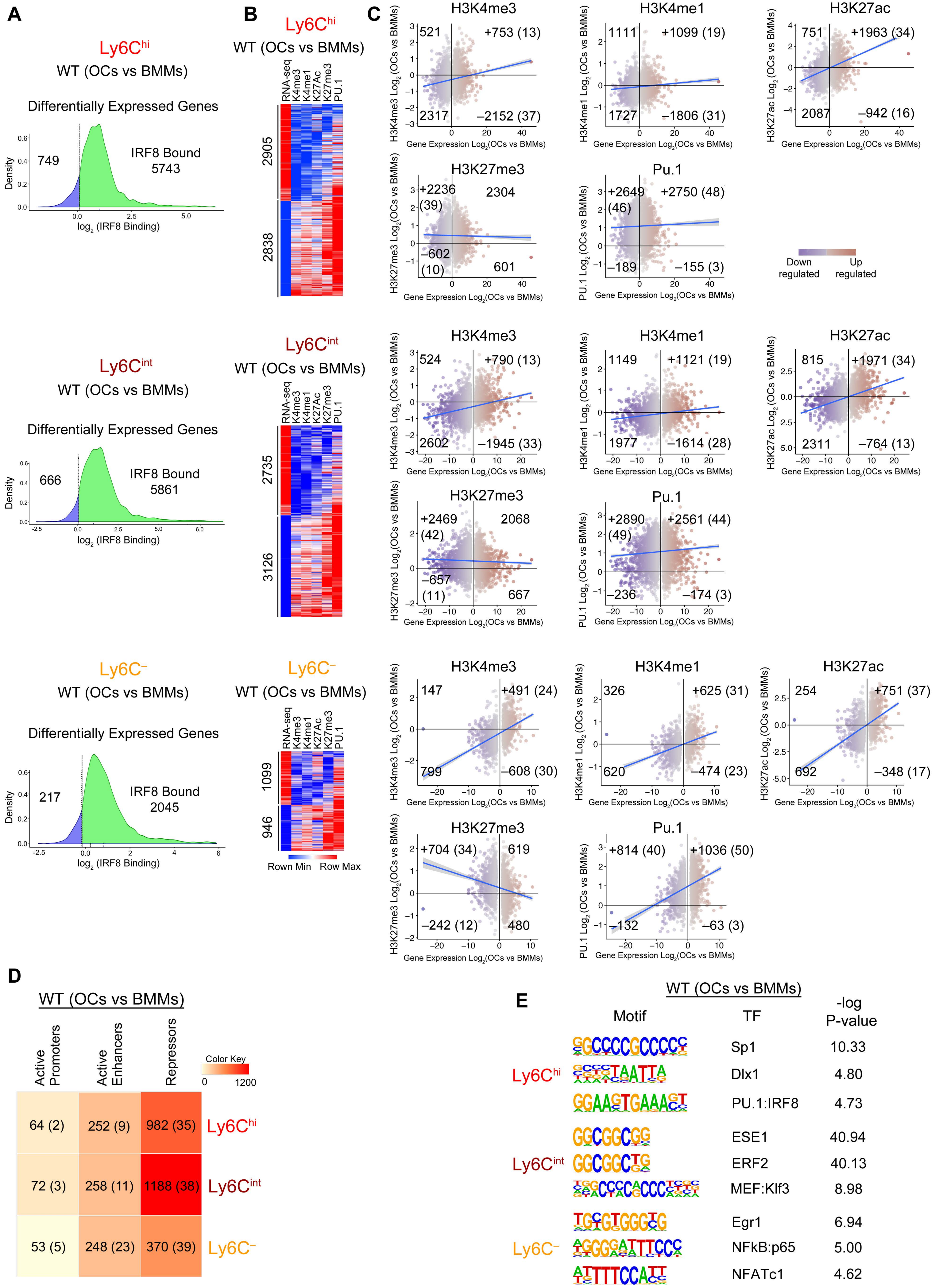

A
Ly6Chi
WT (OCs vs BMMs)
Differentially Expressed Genes
749
IRF8 Bound 5743
Density
log2 (IRF8 Binding)
B
Ly6Chi
WT (OCs vs BMMs)
RNA-seq
K4me3
K4me1
K27Ac
K27me3
PU.1
2905
2838
C
H3K4me3
521
+753 (13)
2317
–2152 (37)
H3K4me1
1111
+1099 (19)
1727
–1806 (31)
H3K27ac
751
+1963 (34)
2087
–942 (16)
H3K27me3
+2236 (39)
2304
–602 (10)
601
Pu.1
+2649 (46)
+2750 (48)
–189
–155 (3)
Gene Expression Log2(OCs vs BMMs)
Down regulated
Up regulated
Ly6Cint
WT (OCs vs BMMs)
Differentially Expressed Genes
666
IRF8 Bound 5861
2735
3126
H3K4me3
524
+790 (13)
2602
–1945 (33)
H3K4me1
1149
+1121 (19)
1977
–1614 (28)
H3K27ac
815
+1971 (34)
2311
–764 (13)
H3K27me3
+2469 (42)
2068
–657 (11)
667
Pu.1
+2890 (49)
+2561 (44)
–236
–174 (3)
Ly6C–
WT (OCs vs BMMs)
Differentially Expressed Genes
217
IRF8 Bound 2045
1099
946
Rown Min
Row Max
H3K4me3
147
+491 (24)
799
–608 (30)
H3K4me1
326
+625 (31)
620
–474 (23)
H3K27ac
254
+751 (37)
692
–348 (17)
H3K27me3
+704 (34)
619
–242 (12)
480
Pu.1
+814 (40)
+1036 (50)
–132
–63 (3)
D
WT (OCs vs BMMs)
Active Promoters
Active Enhancers
Repressors
Color Key
0
1200
64 (2)
252 (9)
982 (35)
Ly6Chi
72 (3)
258 (11)
1188 (38)
Ly6Cint
53 (5)
248 (23)
370 (39)
Ly6C–
E
WT (OCs vs BMMs)
Motif
TF
-log P-value
Ly6Chi
Sp1
10.33
Dlx1
4.80
PU.1:IRF8
4.73
Ly6Cint
ESE1
40.94
ERF2
40.13
MEF:Klf3
8.98
Ly6C–
Egr1
6.94
NFkB:p65
5.00
NFATc1
4.62

**Figure S6, related to Figure 6 and Figure 7. Histone Modifications During Osteoclastogenesis in WT Mice**

(A) Tag density profile show IRF8 bound genes among RANKL-responsive genes differentially expressed in WT OCs vs BMMs.

(B) Heatmap illustrating H3K4me3, H3K4me1, H3K27ac, H3K27me3 and PU.1 binding signal in DEGs between WT OCs vs BMMs.

(C) Scatter plots show the correlation between histone modifications and genes either upregulated or downregulated in WT OCs vs BMMs. Gain in histone marks indicated by + and loss of histone marks indicated by –. Data are represented as $\log_2$.

(D) Heatmap depicting number of active promoters, active enhancers, and repressors acquired in WT OCs vs BMMs. Numbers in parenthesis indicate percentages.

(E) Homer motif analysis of active promoter and active enhancer regions enriched in WT OCs vs BMMs.

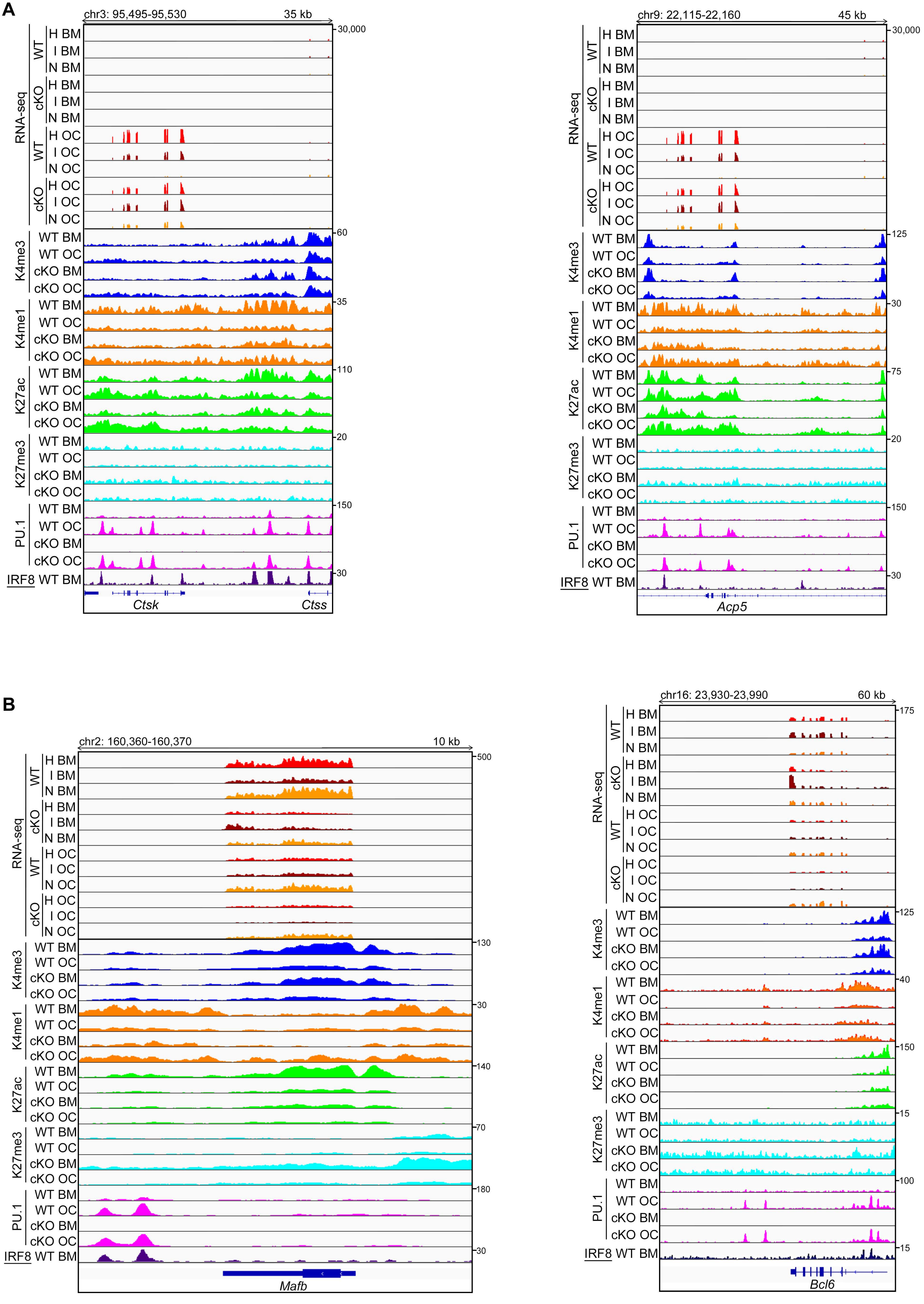

A
chr3: 95,495-95,530
35 kb
RNA-seq
WT
H BM
I BM
N BM
cKO
H BM
I BM
N BM
WT
H OC
I OC
N OC
cKO
H OC
I OC
N OC
30,000
K4me3
WT BM
WT OC
cKO BM
cKO OC
60
K4me1
35
K27ac
110
K27me3
20
PU.1
150
IRF8 WT BM
30
Ctsk
Ctss
chr9: 22,115-22,160
45 kb
30,000
125
30
75
20
150
30
Acp5
B
chr2: 160,360-160,370
10 kb
500
130
30
140
70
180
30
Mafb
chr16: 23,930-23,990
60 kb
175
125
40
150
15
100
15
Bcl6

**Figure S7, related to Figure 6 and Figure 7. The Epigenetic Landscape of Positive and Negative Regulators of Osteoclast Differentiation**

Representative UCSC Genome Browser tracks displaying normalized tag-density profiles at key OC-specific genes.

(A) Note the enrichment of H3K4me1 and H3K27ac marks at the *Ctsk* and *Acp5* loci (positive regulators) in *Irf8 cKO* OCs when compared to WT OCs.

(B) Note the enrichment of H3K27me3 marks at the *Mafb* and *Blc6* loci (negative regulators) in *Irf8 cKO* BMMs and OCs when compared to respective WT cells.
