## Supplementary material for "Monocyte Subsets with High Osteoclastogenic Potential and Their Epigenetic Regulation Orchestrated by IRF8": Key Resource Table

**KEY RESOURCES TABLE**

| **REAGENT or RESOURCE** | **SOURCE** | **IDENTIFIER** |
| --- | --- | --- |
| **Antibodies** |  |  |
| 7-AAD | BioLegend | Cat# 420403 |
| Anti-CD115 APC (eBio) (clone AFS98) | eBioscience | Cat# 17115282 |
| Anti-CD117 PE-Cy7 (clone 2B8) | BioLegend | Cat# 105813 |
| Anti-CD11b Brilliant Violet 421(clone M1/70) | BioLegend | Cat# 101236 |
| Anti-CD11b APC-Cy7 (clone M1/70) | BioLegend | Cat# 101226 |
| Anti-CD11c AF532 (clone N418) | eBioscience | Cat# 58011480 |
| Anti-CD135 (Flt-3) PE-Cy5 (clone A2F10) | BioLegend | Cat# 135312 |
| Anti-CD19 PE (clone 6D5) | BioLegend | Cat# 115507 |
| Anti-CD19 Alexa Fluor 488 (clone CD5) | BioLegend | Cat# 115521 |
| Anti-CD3 PE (clone 17A2) | BioLegend | Cat# 100205 |
| Anti-CD3 APC (clone 17A2) | BioLegend | Cat# 100235 |
| Anti-CD45 PE (clone I3/2.3) | BioLegend | Cat# 147712 |
| Anti-CD45 Alexa Fluor 700 (clone 30-F11) | eBioscience | Cat# 56045180 |
| Anti-CD45R (B220) PE (clone RA3-6B2) | BioLegend | Cat# 103208 |
| Anti-CD45R (B220) Alexa Fluor 488 (clone RA3-6B2) | eBioscience | Cat# 53045280 |
| Anti-IRF8 PerCP-eFluor 710 (clone V3GYWCH) | eBioscience | Cat# 46985280 |
| Anti-Ly-6C eFluor 450 (clone HK1.4) | eBioscience | Cat# 48593282 |
| Anti-Ly-6G PE-Cy7 (clone 1A8) | BioLegend | Cat# 127618 |
| Anti-Ly-6G PE (clone 1A8) | BD Biosciences | Cat# 561104 |
| Anti-MHC II (I-Ab) Alexa Fluor 488 (clone AF6 120.1) | BioLegend | Cat# 116410 |
| Anti-PDCA-1 APC (clone ebio927) | eBioscience | Cat# 17317280 |
| Anti-SiglecH PE-Cy7 (clone ebio440c) | eBioscience | Cat# 25033380 |
| Anti-TER-119 PE (clone TER-119) | BioLegend | Cat# 116208 |
| Anti-TER-119 PE-Dazzle 594 (clone TER-119) | BioLegend | Cat# 116243 |
| Anti-biotin Microbeads Anti-TER-119 PE-Dazzle 594 (clone TER-119) | Miltenyi | Cat# 130100629 |
| Anti-DC-STAMP antibody | Millipore | Cat# MABF39-I |
| Anti-NFAT2 antibody | Abcam | Cat# ab25916 |
| Anti-Cathepsin K antibody | Abcam | Cat# ab19027 |
| Anti-IRF8 antibody | ThermoFischer | Cat# 39-8800 |
| Anti-beta Actin antibody | Abcam | Cat# ab8226 |
| Anti-Rabbit IgG H&L antibody | Abcam | Cat# ab6721 |
| Anti-Mouse IgG H&L antibody | Abcam | Cat# ab6728 |
| Anti-H3K27ac antibody | Abcam | Cat# ab4729 |
| Anti-Histone H3 (mono methyl K4) antibody | Abcam | Cat# ab8895 |
| Anti-Histone H3 (tri methyl K4) antibody | Abcam | Cat# ab8580 |
| Anti-Histone H3 (tri methyl K27) antibody | Millipore | Cat# 07-449 |
| Anti-PU.1 antibody | Laboratory of Michael Ostrowski | (Carey et al., 2018) |
| **Chemicals, Peptides, and Recombinant Proteins** | | |
| Dynabeads Oligo(dT)25 | ThermoFischer | Cat# 61105 |
| Dynabeads Protein G | ThermoFischer | Cat# 10004D |
| cOmplete, EDTA-free Protease Inhibitor Cocktail | Sigma-Aldrich | Cat# 11873580001 |
| Proteinase K Solution | ThermoFischer | Cat# AM2546 |
| High-Capacity cDNA Reverse Transcription Kit | ThermoFischer | Cat# 4374967 |
| SYBR Green PCR Master Mix | ThermoFischer | Cat# 4309155 |
| TRIzol Reagent | ThermoFischer | Cat# 15596026 |
| ACK Lysing Buffer | Quality Biological | Cat# 118-156-101 |
| Halt Protease Inhibitor Cocktail, EDTA-Free | ThermoFischer | Cat# 78425 |
| Recombinant Murine M-CSF | R&D systems | Cat# 416-ML/CF |
| Recombinant Murine RANKL | R&D systems | Cat# 462-TEC/CF |
| **Critical Commercial Assays** | | |
| NEBNext Poly(A) mRNA Magnetic Isolation kit | NEB | Cat# E7490 |
| NEBNext Ultra II Directional RNA Library Prep kit | NEB | Cat# E7760 |
| NEBNext Multiplex oligos for Illumina | NEB | Cat# E7335 |
| NEBNext Ultra II DNA library preparation kit | NEB | Cat# E7645 |
| QIAquick PCR Purification Kit | Qiagen | Cat# 28104 |
| RNeasy Mini Kit | Qiagen | Cat# 74104 |
| Monocyte Isolation Kit (BM), mouse | Miltenyi | Cat# 130-100-629 |
| RatLaps (CTX-I) EIA | immunodiagnostic systems | Cat# AC-06F1 |
| High Sensitivity DNA ScreenTape Analysis (D1000) | Agilent | Cat# 5067-5587 |
| **Deposited Data** | | |
| RNA-seq and ChIP-seq | GEO | GSE151483 |
| **Experimental models: Organisms/Strains** | | |
| Mouse: *Irf8^fl/fl^* | Laboratory of Keiko Ozato and Herbert Morse | (Feng et al., 2011) |
| Mouse: *Csf1r^cre^* | Jackson Laboratory | Jax Stock# 021024;  (Deng et al., 2010) |
| Mouse: *Irf8 gKO* and *Irf8 gWT* | Laboratory of Keiko Ozato and Herbert Morse | (Tamura and Ozato, 2002; Thumbigere-Math et al., 2019) |
| Mouse: *Irf8 cWT* (*Irf8^fl^; Csf1r^cre/-^*) | Generated in this study | N/A |
| Mouse: *Irf8 cKO* (*Irf8^fl/fl^; Csf1r^cre/+^*) | Generated in this study | N/A |
| **Oligonucleotides** | | |
| Primers for qPCR, see Table S1 | This paper | N/A |
| **Software and Algorithms** | | |
| GraphPad Prism 8 (version 8.0.1) | GraphPad Software, Inc., Carlifornia | <https://www.graphpad.com> |
| ImageJ | ImageJ | <https://imagej.nih.gov/ij/> |
| AnalyzePro 1.0 | AnalyzeDirect | <https://analyzedirect.com> |
| FCS Express 6 | De Novo Software | <https://denovosoftware.com> |
| Bowtie2 aligner version 2.4.0 | Langmead and Salzberg, 2012 | <http://bowtie-bio.sourceforge.net/bowtie2/index.shtml> |
| HOMER | Heinz et al., 2010 | <http://homer.ucsd.edu/homer/introduction/install.html> |
| GSEA | Broad Institute | <https://www.gsea-msigdb.org/gsea/index.jsp> |
| R package: DESeq2 | Anders and Huber, 2010 | <https://bioconductor.org/packages/release/bioc/html/DESeq2.html> |
| Ingenuity Pathway Analysis | N/A | <https://www.qiagenbioinformatics.com/products/ingenuity-pathway-analysis/> |
| **Other** | | |
| Cytek Aurora | Cytek Biosciences | N/A |
| BD FACS Aria II cell sorter | BD Biosciences | N/A |
| Zeiss Axio Observer 3 Microscope | Carl Zeiss | N/A |
| RNA-sequencing | Genome Technology Unit, NIAMS/NIH | N/A |
| ChIP-sequencing | Genome Technology Unit, NIAMS/NIH | N/A |
