## Supplemental Table 1 for "Monocyte Subsets with High Osteoclastogenic Potential and Their Epigenetic Regulation Orchestrated by IRF8"

| Mouse *Hprt* | Forward | 5' - TCAGTCAACGGGGGACATAAA - 3' |
| --- | --- | --- |
|  | Reverse | 5' – GGGGCTGTACTGCTTAACCAG - 3' |
| Mouse *Nfatc1* | Forward | 5' – TCATCCTGTCCAACACCAAA - 3' |
|  | Reverse | 5' – TCACCCTGGTGTTCTTCCTC - 3' |
| Mouse *Ctsk* | Forward | 5' – AGGGAAGCAAGCACTGGATA - 3' |
|  | Reverse | 5' – GCTGGCTGGAATCACATCTT - 3' |
| Mouse *Oscar* | Forward | 5' – TCATCTGCTTGGGCATCATA - 3' |
|  | Reverse | 5' – ACAAGCCTGACAGTGTGGTG - 3' |
| Mouse *Acp5* | Forward | 5' – CGTCTCTGCACAGATTGC - 3' |
|  | Reverse | 5' – GAGTTGCCACACAGCATCAC - 3' |
| Mouse *Dcstamp* | Forward | 5' – GGGCACCAGTATTTTCCTGA - 3' |
|  | Reverse | 5' – TGGCAGGATCCAGTAAAAGG - 3' |
| Mouse *Irf8* | Forward | 5' – GATCGAACAGATCGACAGCA - 3' |
|  | Reverse | 5' – GCTGGTTCAGCTTTGTCTCC - 3' |
| Mouse *Csf1r* | Forward | 5' – TGTCATCGAGCCTAGTGGC - 3' |
|  | Reverse | 5' – GGTCCAAGGTCCAGTAGGG - 3' |

**Table S1 qPCR primers, related to Figures 2 and 4, and Figures S1 and S5**
